# A physical mechanism of TANGO1-mediated bulky cargo export

**DOI:** 10.1101/576637

**Authors:** Ishier Raote, Morgan Chabanon, Nikhil Walani, Marino Arroyo, Maria F. Garcia-Parajo, Vivek Malhotra, Felix Campelo

## Abstract

The endoplasmic reticulum (ER)-resident transmembrane protein TANGO1 assembles into a ring around COPII subunits at ER exit sites (ERES), and links cytosolic membrane-remodeling machinery, tethers, and ER-Golgi intermediate compartment (ERGIC) membranes to procollagens in the ER lumen (Raote *et al*., 2018). This arrangement is proposed to create a route for direct transfer of procollagens from ERES to ERGIC membranes via a tunnel. Here, we present a physical model in which TANGO1 forms a linear filament that wraps around COPII lattices at ERES to stabilize the neck of a growing transport intermediate. Importantly, our results show that TANGO1 is able to stabilize ERES-ERGIC opening by regulating ER membrane tension to allow procollagen loading and export from the ER. Altogether, our theoretical approach provides a mechanical framework of TANGO1 as a membrane tension regulator to control procollagen export from the ER.

## INTRODUCTION

Multicellularity requires not only the secretion of signaling proteins –such as neurotransmitters, cytokines, and hormones– to regulate cell-to-cell communication, but also of biomechanical matrices composed primarily of proteins such as collagens, which form the extracellular matrix (ECM) (Kadler *et al*., 2007; Mouw, Ou and Weaver, 2014). These extracellular assemblies of collagens are necessary for tissue biogenesis and maintenance. Collagens, like all conventionally secreted proteins, contain a signal sequence that targets their entry into the endoplasmic reticulum (ER). After their glycosylation, procollagens fold and trimerize into a characteristic triple-helical structure, which is supposed to be very rigid and long (McCaughey and Stephens, 2019). These bulky procollagens are then exported from the ER at specialized export domains, termed ER exit sites (ERES). ERES are a fascinating subdomain of the ER, but a basic understanding of how these domains are created and segregated from the rest of the ER for the purpose of cargo export still remains a challenge. The discovery of TANGO1 as a key ERES-resident player, has made the processes of procollagen export and the organization of ERES amenable to molecular analysis (Bard *et al*., 2006; Saito *et al*., 2009; Wilson *et al*., 2011).

In the lumen of the ER, the SH3-like domain of TANGO1 binds procollagen via HSP47 (Saito *et al*., 2009; Ishikawa *et al*., 2016) (***Figure 1A***). On the cytoplasmic side, TANGO1 has a proline-rich domain (PRD) and two coiled-coil domains (CC1 and CC2) (***Figure 1A***). The PRD of TANGO1 interacts with the COPII components Sec23A and Sec16 (Saito *et al*., 2009; Ma and Goldberg, 2016; Maeda, Katada and Saito, 2017); the CC1 domain binds the NBAS/RINT1/ZW10 (NRZ) tethering complex to recruit ER-Golgi intermediate compartment (ERGIC) membranes and also drives self-association amongst TANGO1 proteins (Santos *et al*., 2015; Raote *et al*., 2018); and the CC2 domain oligomerizes with proteins of the TANGO1 family (such as cTAGE5) (Saito *et al*., 2011; Maeda, Saito and Katada, 2016). Both cytosolic and lumenal activities of TANGO1 are critical for its function. A recent report identified a disease-causing mutation in TANGO1 in a human family, which results in a substantial fraction of TANGO1 protein being truncated lacking its cytosolic functions leading to various skeletal abnormalities and collagen export defects (Lekszas *et al*., 2020). Recently, we visualized procollagen export domains with high lateral spatial resolution using stimulated emission depletion (STED) nanoscopy in mammalian tissue cultured cells (Raote *et al*., 2017, 2018). These studies revealed that TANGO1 organizes at the ERES into ring-like structures, of ~200 nm in lumenal diameter, that corral COPII components. Moreover, two independent studies showed that TANGO1 rings are also present in *Drosophila melanogaster* (Liu *et al*., 2017; Reynolds *et al*., 2019).

**Figure 1.**
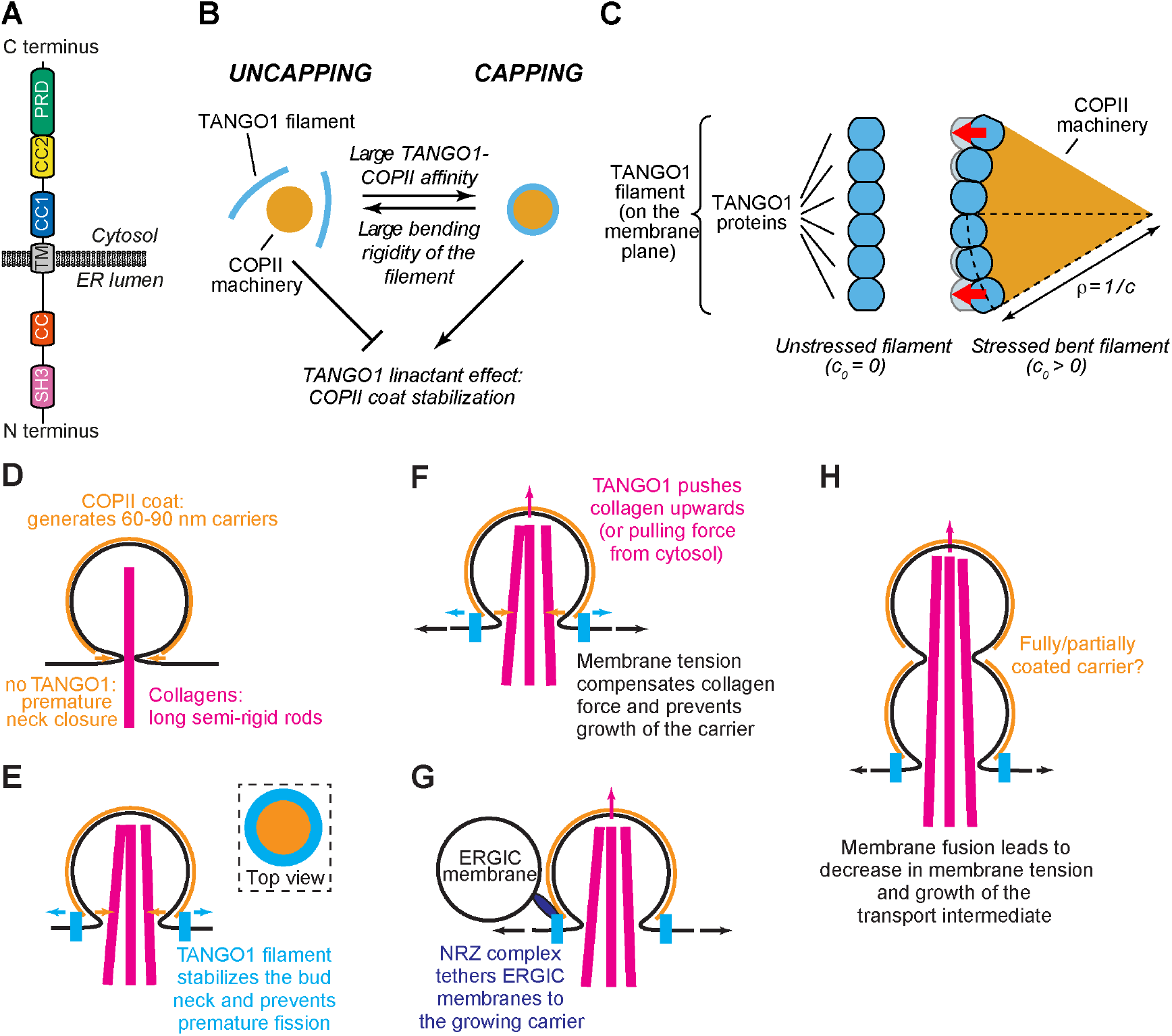
Physical model of TANGO1-dependent transport intermediate formation. **(A)** Schematic representation of the domain structure and topology of TANGO1, indicating the SH3 domain, a luminal coiled-coiled domain (CC), the one and a half transmembrane region (TM), the coiled-coiled 1 (CC1) and 2 (CC2) domains, and the PRD. **(B)** Schematic description of the TANGO1 ring formation model. ERES consisting of COPII subunits assemble into in-plane circular lattices (orange), whereas proteins of the TANGO1 family assemble into filaments by lateral protein-protein interactions (light blue). A tug-of-war between the affinity of the TANGO1 filament to bind COPII subunits (promoting capping of peripheral COPII subunits) and the resistance of the filament to be bent (promoting uncapping) controls the capping-uncapping transition. Only when TANGO1 caps the COPII lattice, it acts as a linactant by stabilizing the peripheral COPII subunits. **(C)** Schematic representation of individual proteins that constitute a TANGO1 filament and of how filament bending is associated with elastic stress generation. Individual TANGO1 family proteins (blue shapes) bind each other in a way that is controlled by the structure of protein-protein binding interfaces, leading to formation of an unstressed filament of a certain preferred curvature, *c*_0_ (left cartoon, showing the case where *c*_0_ = 0). Filament bending can be caused by the capping of TANGO1 filament to peripheral COPII machinery (orange area), which generates a stressed bent filament of a certain radius of curvature, *R* = 1/*c* (right cartoon). Such deviations from the preferred shape (shown in light blue) are associated with elastic stress generation (red arrows point to the direction of the filament reaction forces to the generated stresses, which correspond to the direction of recovery of the filament preferred shape). **(D)** In the absence of functional TANGO1, COPII coated spherical vesicles assemble normally, generating spherical carriers of between 60–90 nm in size. Procollagens cannot be packed into such small carriers. **(E)** A TANGO1 filament siting at the base of a growing COPII patch encircles COPII components as experimentally observed (see top view in the top right subpanel) and packages procollagens to the export sites. This TANGO1 fence can serve to stabilize the neck of the transport carrier hence preventing the premature formation of a small carrier. **(F)** A possible cytosolically-directed force (procollagen pushing from the inside or a pulling force from the cytosol) can work in the direction of generating a large intermediate. By contrast, large membrane tensions work to prevent carrier elongation. **(G)** The NRZ complex (dark blue), which is recruited to the procollagen export sites by the TANGO1 TEER domain, tethers ERGIC53-containing membranes. **(H)** Fusion of these tethered membranes can lead to a local and transient decrease in the membrane tension, which can allow for the growth of the transport intermediate to be able to include the long semi-rigid procollagen molecules. Whether the intermediate is fully or only partially coated still remains unknown.

To further extend these findings, we combined STED nanoscopy with genetic manipulations and established that TANGO1 rings are organized by *(i)* lateral self-interactions amongst TANGO1-like proteins, *(ii)* radial interactions with COPII subunits, and *(iii)* tethering of small ER-Golgi intermediate compartment (ERGIC) vesicles to assist in the formation of a procollagen-containing export intermediate (Raote *et al*., 2018). Overall, the data suggest a mechanism whereby TANGO1 assembles into a ring, which selectively gathers and organizes procollagen, remodels COPII budding machinery, and recruits ERGIC membranes for the formation of a procollagen-containing transport intermediate. However, the biophysical mechanisms governing these events and how they are regulated by TANGO1 remain unknown. Here, we present and analyze a biophysical model of TANGO1 ring assembly around polymerizing COPII-coated structures for procollagen export. Our model allows us to address: *(i)* the physical mechanisms by which TANGO1 and its interactors can assemble into functional rings at ERES, forming a fence around COPII coat components; and *(ii)* whether and how a TANGO1 fence can couple COPII polymerization and regulate membrane tension to create an export route for procollagens at the ERES. Overall, we propose a novel mechanism of TANGO1-regulated procollagen export, which consists of two sequential steps. First, TANGO1 rings, at the edge of a polymerizing COPII structure, stabilize the neck of a growing procollagen-containing export intermediate and thus prevent premature fission. Second, carrier growth can be stimulated by the ability of TANGO1 to act as a membrane tension regulator by tethering and fusing ERGIC membranes. Importantly, we show that TANGO1-mediated local reduction of the membrane tension at the ERES changes the free energy profile of the system to promote carrier growth.

## PHYSICAL MODEL OF TANGO1-ASSISTED TRANSPORT INTERMEDIATE FORMATION

Can TANGO1 modulate the shape of a growing bud to accommodate large and complex cargoes? And, if so, would a TANGO1 ring structure be especially suited to achieve this task? To answer these questions, we assembled a physical model of transport intermediate formation that incorporates the effects of TANGO1 ring formation. In our model, we consider different scenarios under which TANGO1 can modulate the shape of COPII-dependent carriers. We first describe how TANGO1 rings can form around COPII patches and then propose a general model of carrier formation that includes the contribution of TANGO1 rings.

### QUALITATIVE DESCRIPTION OF TANGO1 RING ASSEMBLY

To assess and rationalize the assembly of TANGO1 into rings at ERES, we propose a physical model built on the accumulated experimental data. First, we hypothesize that TANGO1 forms a filament that can be held together by lateral protein-protein interactions between TANGO1-family proteins (TANGO1, cTAGE5 and TANGO1-Short) (Raote *et al*., 2018). This hypothesis is based on the following observations: *(i)* TANGO1 is seen in ring-like filamentous assemblies by STED nanoscopy (Raote *et al*., 2017); *(ii)* there is a direct 1:1 binding between TANGO1 and cTAGE5 CC2 domains (Saito *et al*., 2011); *(iii)* TANGO1-Short and cTAGE5 can form oligomers and oligomeric complexes together with Sec12 and TANGO1 (Maeda, Saito and Katada, 2016); *(iv)* TANGO1 and TANGO1-Short can directly homo-dimerize by their CC1 domains (Raote *et al*., 2018); and *(v)* super-resolution live lattice SIM imaging of TANGO1 in the *D. melanogaster* larval salivary gland shows filament growth in ring formation (Reynolds *et al*., 2019). Such a filament forms by the assembly of TANGO1-family proteins, which we propose occurs in a linear or quasi-linear fashion, rather than as a protein aggregate or protein cluster. Depending on the structural details of the interactions between TANGO1 proteins, the filament would tend to adopt a defined shape, and deviations from such a shape would necessitate the supply of external energy. In terms of elasticity of the filament, such a filament is subject to internal strains and stresses and therefore resists bending away from its preferred shape or curvature. Evidence for the existence of linear assemblies of transmembrane proteins has indeed been reported in the context of transmembrane actin-associated (TAN) lines that couple outer nuclear membrane components to actin cables (Luxton *et al*., 2010).

Second, TANGO1 binds the inner layer of the COPII coat (Saito *et al*., 2009). We hypothesize that TANGO1 stabilizes the edges of the COPII lattice by binding to peripheral COPII subunits, thereby effectively reducing COPII lattice line energy at the ERES (Glick, 2017). COPII coat assembly at the ERES occurs by polymerization of the individual COPII subunits into a lattice (Aridor, 2018; Peotter *et al*., 2019). This process starts with activation and membrane binding of the Sar1 GTPase, which recruits Sec23-Sec24 heterodimers that form the inner layer of the COPII coat. Subsequently, the second layer of the coat, composed of Sec13-Sec31 subunits, is recruited to the ERES, eventually leading to the budding of a COPII-coated vesicle. The free energy of coat polymerization includes the binding free energy of the COPII subunits, the elastic penalty of bending the membrane underneath, and also the line energy due to the unsatisfied binding sites of COPII subunits occupying the edges of the growing lattice (Saleem *et al*., 2015). Because proteins of the TANGO1 family physically interact with the COPII components Sec23, Sec16, and Sec12, we hypothesize that by binding to COPII subunits placed at the periphery of the growing coat (Ma and Goldberg, 2016; Hutchings *et al*., 2018; Raote *et al*., 2018), TANGO1 stabilizes the domain boundary, effectively reducing its line energy. In analogy to surfactants –molecules that adsorb into liquid-liquid two-dimensional interfaces decreasing their surface tension–, we propose that by binding to COPII subunits, TANGO1 proteins act as line-active agents, or *linactants* (Trabelsi *et al*., 2008). In the context of HIV gp41-mediated membrane fusion, linactant compounds, such as vitamin E, lower the interfacial line tension between different membrane domains to inhibit HIV fusion (Yang, Kiessling and Tamm, 2016).

Qualitatively, our model for TANGO1 ring assembly can be described as a tug-of-war between two different driving forces: the resistance to bending of TANGO1 filaments –driven by the nature of the TANGO1-TANGO1 protein interactions–, and the binding affinity of TANGO1 proteins for peripheral COPII subunits. These different forces can control the formation of TANGO1 rings around COPII coats at ERES at different stages of transport intermediate formation. For instance, if the resistance to bending of the TANGO1 filament is relatively small or the binding affinity of TANGO1 for the COPII subunits is relatively large, the filament will easily wrap around COPII patches, forming a TANGO1 ring (a process we refer to as COPII wetting or *capping*) (***Figure 1B,C***). As a result, the linactant effect of TANGO1 on COPII-coated ERES that will reduce the line energy, thus limiting further growth of the COPII lattice and the size of the TANGO1 rings (***Figure 1B***). By contrast, if TANGO1 filaments are very rigid or the affinity of TANGO1 proteins for COPII subunits is low (for instance, in cells expressing mutants of TANGO1 with reduced or abrogated interaction to COPII proteins), COPII capping by the filament will be energetically unfavorable and as a result, TANGO1 will not act as a COPII linactant (***Figure 1B***).

### TANGO1-DEPENDENT TRANSPORT INTERMEDIATE FORMATION

The formation of canonical coated transport carriers (such as COPI-, COPII-, or clathrin-coated carriers) relies on the polymerization on the membrane surface of a large-scale protein structure: the protein coat. Polymerized coats usually adopt spherical shapes, which bend the membrane underneath accordingly (Faini *et al*., 2013), although helical arrangements of COPII coats have been proposed (Zanetti *et al*., 2013; Ma and Goldberg, 2016). Coat polymerization promotes bending of the underlying membrane if the binding energy of the coat to the membrane is larger than the energy required to bend the membrane and if the coat structure is more rigid than the membrane (Derganc, Antonny and Čopič, 2013; Kozlov *et al*., 2014; Saleem *et al*., 2015). Hence, in the absence of a functional TANGO1, COPII coats generate standard 60-90 nm spherical transport carriers (***Figure 1D***). In this situation, the neck of the growing carrier prematurely closes without being able to fully incorporate long semi-rigid procollagen molecules, which are either not fully packaged or not efficiently recruited to the COPII export sites due to the lack of TANGO1 (***Figure 1D***). In our model for TANGO1 ring formation, we proposed that one of the potential roles of such a ring is to act as a linactant to stabilize free COPII subunits at the edge of the polymerized structure (Glick, 2017; Raote *et al*., 2017) and hence prevent or kinetically delay the premature closure of the bud neck (***Figure 1E***). Moreover, mechanical forces directed towards the cytosolic side of the bud, either from the ER lumen (e.g. TANGO1 pushing procollagen upwards) or from the cytosol (e.g. molecular motors pulling on the growing bud), will induce growth of the transport intermediate (Derényi, Jülicher and Prost, 2002; Roux *et al*., 2002; Koster *et al*., 2003; Leduc *et al*., 2004; Watson *et al*., 2005; Pinot, Goud and Manneville, 2010) (***Figure 1F***). Such pulling forces can be counterbalanced by membrane tension, which generally acts as an inhibitory factor preventing bud formation (Saleem *et al*., 2015; Hassinger *et al*., 2017; Wu *et al*., 2017) (***Figure 1F***). In parallel, TANGO1 recruits the NRZ complex that tethers ERGIC53-containing membranes in apposition to TANGO1 rings (Raote *et al*., 2018) (***Figure 1G***). Fusion of such vesiculo-tubular structures to the budding site would deliver membrane lipids to the ER membrane, which rapidly and transiently drops local membrane tension, hence overcoming the tension-induced arrest in transport intermediate growth (***Figure 1H***). Mixing of membrane components is prevented by the particular structure of TANGO1 transmembrane helices, which create diffusion barrier (Raote *et al*., 2020). The presence of long and rigid procollagen molecules (Buehler, 2008; McCaughey and Stephens, 2019) in the lumen of the immature carrier could sterically hinder the full closure of the necks between the subsequent spherical pearls, thus contributing to fission inhibition and enlargement of the transport intermediate (Peotter *et al*., 2019) (***Figure 1H***). The shape and coat coverage of procollagen-containing export intermediates remain, to the best of our knowledge, a matter of speculation. Both large pearled tubes (***Figure 1H***) or cylindrical vesicles have been proposed to function at the level of the ER membrane (Bannykh, Rowe and Balch, 1996; Mironov *et al*., 2003; Watson and Stephens, 2005; Zeuschner *et al*., 2006; Robinson *et al*., 2015; Gorur *et al*., 2017; Omari *et al*., 2018; Yuan *et al*., 2018). Remarkably, multibudded, pearled-like structures have been observed in a COPII *in vitro* budding system (Bacia *et al*., 2011). We recently proposed the alternative possibility that a short-lived, transient direct tunnel between the ER and the ERGIC/Golgi complex can allow for the directional export of cargoes from the ER (Raote and Malhotra, 2019). In our model, TANGO1 rings help prevent the fission of the carrier and thus allow for the formation of such tunnels between the ER and the ERGIC. Finally, although there is experimental evidence of tubular COPII polymerization in vitro as observed by cryo-electron tomography (Zanetti *et al*., 2013; Hutchings *et al*., 2018), COPII coats have a preference to polymerize into spherical structures. Hence, for the sake of clarity we will base our analytical model of membrane deformation by COPII coats on the assumption that COPII polymerization on the membrane imposes a spherical shape of fixed curvature to the underlying membrane. A more general model where the condition of imposed spherical curvature is relaxed is presented in ***Appendix 1***.

To quantitate the feasibility of the proposed pathway of transport intermediate growth (***Figure 1D–H***), we developed a physical model that accounts for the relative contribution of each of the aforementioned factors to the overall free energy of the system. Such a model allows us to predict *(i)* the shape transitions from a planar membrane to incomplete buds and to large transport intermediates; and *(ii)* the formation of TANGO1 rings around COPII components, which we refer to as capping of COPII by TANGO1. Intuitively, one can see that COPII polymerization favors the formation of spherical buds, whereas TANGO1 linactant strength and filament bending prevent neck closure by capping incomplete COPII buds. Outward-directed forces promote the growth of large intermediates, whereas large membrane tension inhibits such a growth. Taking advantage of a recently developed theoretical model of membrane elasticity in the context of clathrin-coated vesicle formation (Saleem *et al*., 2015), we expand on this model to include the contributions of TANGO1-like proteins in modulating COPII-dependent carrier formation, the description of which follows.

We consider that the ER membrane is under a certain lateral tension, *σ*, and resists bending with a bending rigidity, *κ_b_*. Growth of a COPII bud starts by COPII polymerization into a spherical shape of radius *R*, driven by the assembly of individual subunits. In physical terms, lattice growth is favored if the free energy of the system decreases upon assembly of soluble subunits into the lattice. Hence, the chemical potential of the polymerizing COPII coat, *μ_c_*, includes the COPII binding chemical potential, *μ_c_*^0^, and the term associated with the bending energy of the underlying membrane (see Materials and Methods and (Saleem *et al*., 2015)). Energetically favorable lattice polymerization is assisted by the relatively large binding energies between COPII subunits, which in turn penalize the existence of free binding sites within the COPII lattice. In particular, the peripheral subunits laying at the edge of the lattice have a number of unsatisfied bonds, which implies that the free energy of the system could be further relaxed by fulfilling those bonds and hence reducing the length of the COPII patch edge. This is formalized in physical terms by considering a line energy of the COPII lattice, which is proportional to its boundary length, *l* = 2*πρ*, and the proportionality factor is the line tension of the free, peripheral COPII subunits, *λ*_0_. Altogether, these contributions give the part of the free energy per unit area that is independent of TANGO1, *f_c_*^0^, which can be written as

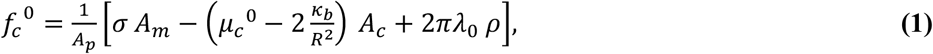

where *A_m_* is the membrane surface area, *A_c_* is the surface area of the membrane covered by the COPII coat, *A_p_* is the surface area of the carrier projection onto the flat membrane, and *ρ* is the radius of the base of the carrier (see ***Figure 1–figure supplement 1*** and Materials and methods section).

To quantitatively analyze the effect of TANGO1 ring formation and COPII capping on transport intermediate formation, we use a continuum model, which implicitly considers TANGO1 family proteins (TANGO1, cTAGE5 and TANGO1-Short) and TANGO1-binding COPII subunits. Here, the “microscopic” interaction energies are averaged out into “macroscopic” free energies, such as the filament bending energy, or the coat line energy. In addition, molecular length scales are represented by effective quantitates, such as the spontaneous curvature of the rings or the imposed coat radius of curvature. Although simplistic in nature, this continuum model is a suitable choice for a semi-quantitative description of the main physical mechanisms driving ring formation, as structural data on TANGO1 proteins are currently lacking. Hence, we need to consider two different protein interactions: *(i)* TANGO1-TANGO1 interactions, which control the bending energy of the TANGO1 filament; and *(ii)* TANGO1 interaction with peripheral COPII subunits, which controls the line energy of the COPII domain. To build our physical model of TANGO1 ring formation, we need a mathematical formulation of these interactions in terms of physical energies.

#### Bending energy of the TANGO1 filament

We hypothesized that TANGO1 family proteins have the ability to form linear filaments by protein-protein interactions (***Figure 1C***). Filament deformations generate elastic stresses that arise as a result of filament bending away from its preferred shape. We can use the elastic theory of rods (Landau and Lifshitz, 1986) to describe the bending energy of a TANGO1 filament per unit length as 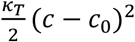, where *κ_T_* is an elastic parameter that characterizes the bending rigidity of the TANGO1 filament, *c* = 1/*ρ* and are the local and preferred curvatures of the filament, respectively, and *ρ* is the local radius of curvature of the filament (***Figure 1C***). If we define the capping fraction, ω, as the fraction of COPII lattice boundary length associated with TANGO1 molecules, we can write down the free energy of TANGO1 filament bending per unit area as

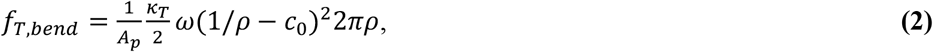

where *ρ* corresponds to the in-plane projected radius of the COPII patch encircled by the TANGO1 ring, and hence also to the radius of curvature of the TANGO1 filament forming the ring (see Materials and Methods).

#### Linactant effect of TANGO1 capping the COPII lattice

We hypothesize that TANGO1 has the ability to cap growing COPII buds by binding to their peripheral subunits. Such TANGO1-COPII binding leads to an effective reduction of the line energy of the bud (see Materials and Methods). The binding affinity of TANGO1 to COPII is Δ*λ*, which can be understood as the linactant strength of TANGO1 capping peripheral COPII subunits. Altogether, we can express the interaction free energy per unit area between TANGO1 and the COPII lattice, *f*_*TANGO*1–*COPII*_, as

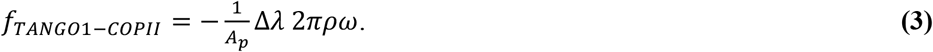

#### Overall free energy of the system

Finally, we also account for the mechanical work of an outward-directed force, *N*, which favors transport intermediate elongation (see Materials and Methods). Altogether, we can write the total free energy per unit surface area, *f_c_*, as 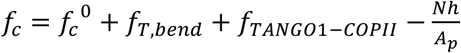, where *h* is the height of the carrier (***Figure 1—figure supplement 1B***). In order to relate the free energy of the system with respect to a reference state, we choose to use the uncoated, flat membrane as the reference state. This state is characterized by the absence of a coat, that is *A_c_* = 0, *λ*_0_ = 0, and ω = 0; the absence of an applied force, *N* = 0; and given that the membrane is flat, we have that *A_m_ = A_p_*, and *h* = 0. Hence, we can write, 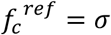, and define the free energy change of the system, 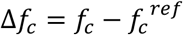, as

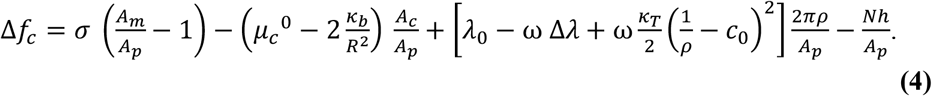

### PARAMETER ESTIMATION

The free energy per unit area, ***Equation (4)*** depends on a number of structural, biochemical, and mechanical parameters, which we can split in three groups: *(i)* membrane-associated parameters, *(ii)* coat-associated parameters, and *(iii)* TANGO1-associated parameters (***Table 1***). As for the membrane-associated parameters, we have the lateral tension, *σ*, and the bending rigidity, *κ_b_*. Regarding membrane-associated parameters, we use the experimentally measured values of the standard membrane tension of the ER, *σ_ER_* = 0.003 *k_B_T/nm*^2^ (Upadhyaya and Sheetz, 2004); and of the membrane bending rigidity, *κ_b_* = 20 *k_B_T* (Niggemann, Kummrow and Helfrich, 1995) (***Table 1***). Regarding coat-associated parameters, we use the size of the standard spherical COPII vesicle, *R* = 37.5 *nm* (Miller and Schekman, 2013). The line tension and the binding free energy of the polymerizing COPII coat, *λ*_0_ and *μ_c_*^0^, respectively, have not been, to the best of our knowledge, experimentally measured. Nevertheless, for clathrin coats, which lead to the formation of vesicles of a size comparable to the standard COPII vesicles, these values have been recently measured, yielding a value of *λ_clathrin_* = 0.05 *pN* for the line tension and of *μ_clathrin_*^0^ = 0.024 ± 0.012 *k_B_T/nm*^2^ for the binding free energy (Saleem *et al*., 2015). We use these values as starting estimations for COPII coats, which we will then vary within a certain range (***Table 1***). Finally, regarding the TANGO1-associated parameters, which are associated to different protein-protein interactions, we have the bending rigidity of the TANGO1 filament, *κ_T_*; the preferred curvature of the filament, *c*_0_; and the TANGO1-COPII binding energy/linactant strength of TANGO1, Δ*λ*. The elastic parameters of the TANGO1 filament, *κ_T_* and *c*_0_, depend on the chemistry of the bonds between the different proteins within a TANGO1 filament. As we lack experimental data on the value of these parameters, we consider them within a wide range of reasonable values. Typical values of the bending rigidity of intracellular filaments formed by protein-protein interactions, such as intermediate filaments, are of the order of *κ_IF_* = 2000 *pN · nm*^2^ (Fletcher and Mullins, 2010), which we consider as an upper limit for the rigidity of a TANGO1 filament. In addition, by taking *κ_T_* = 0, we can exploit our model to study the case where TANGO1 proteins do not form a cohesive filament by attractive lateral protein-protein interactions, but individual proteins can still bind COPII subunits and hence act as monomeric linactants. For our analytical analysis we will start by taking a zero spontaneous curvature of the TANGO1 filament, = 0, and later study it within a range given by twice the radius of experimentally measured TANGO1 rings, −0.02 *nm*^−1^ < *c*_0_ < 0.02 *nm*^−1^. For the value of Δ*λ*, we can make an upper limit estimate, by considering that the TANGO1-COPII binding energy should be lower than the corresponding binding energy between polymerizing COPII components, that is, Δ*λ l*_3_ < *μ_c_*^0^ *l*_1_*l*_2_, where *l*_1_ ≈ 16 *nm* and *l*_2_ ≈ 10 *nm* are the lateral dimensions of the inner COPII coat components Sec23/24 (Matsuoka *et al.*, 2001). Hence, our estimation gives that Δ*λ* < 0.24 *k_B_T/nm*, and therefore we use as the initial value for our analysis half of the upper limit value, Δ*λ* = 0.12 *k_B_T/nm*, which is ten-fold larger than the bare line tension of the coat (***Table 1***).

**TABLE 1:** Parameters used in the large transport intermediate formation model. The free energy ***Equation (4)*** depends on a number of different parameters, which are described in this table.

| Parameter | Description | Value | Notes | Reference |
| --- | --- | --- | --- | --- |
| $\sigma$ | Membrane tension | $0.003 k_B T / nm^2$ (ER);<br>$0.0012 k_B T / nm^2$ (Golgi membranes) | | (Upadhyaya and Sheetz, 2004) |
| $\kappa_b$ | Membrane bending rigidity | $20 k_B T$ | | (Niggemann, Kummrow and Helfrich, 1995) |
| $R$ | Radius of curvature of the COPII coat | $37.5 nm$ | | (Miller and Schekman, 2013) |
| $\lambda_0$ | Bare coat line tension | $0.012 k_B T / nm$ | Not measured for COPII. Used the clathrin value as a reference | (Saleem *et al.*, 2015) |
| $\mu_c^0$ | COPII coat binding energy | $0.024 \pm 0.012 k_B T / nm^2$ | Not measured for COPII. Used the clathrin values as a reference | (Saleem *et al.*, 2015) |
| $\kappa_T$ | TANGO1 filament bending rigidity | $120 k_B T nm$ | Not measured. Range based on standard filament rigidities (see text) | |
| $c_0$ | TANGO1 filament spontaneous curvature | $(-0.02, 0.02) nm^{-1}$ | Not measured. Range based on observed TANGO1 ring sizes. | (Raote *et al.*, 2017) |
| $\Delta\lambda$ | Linactant TANGO1 effect | $0.12 k_B T / nm$ | Not measured. Range based on protein-protein affinity (see text) | - |
| $N$ | Outwards directed force | $0 - 5 k_B T / nm$ | Not measured. Range based on known intracellular forces. | (Kovar and Pollard, 2004) (Actin); (Block *et al.*, 2003) (Molecular motors) |

## RESULTS

### ENERGETICALLY ACCESSIBLE EQUILIBRIUM CONFIGURATIONS

We consider that the carrier adopts the equilibrium configuration, corresponding to the configuration of minimum free energy, ***Equation (4)***. Although the system is not in equilibrium, this assumption will be valid as long as the mechanical equilibration of the membrane shape is faster than the fluxes of the lipids and proteins involved in the problem (Sens and Rao, 2013; Campelo *et al*., 2017). Hence, assuming local equilibrium, our aim is to compute the shape of the carrier and the state of TANGO1 capping that minimize ***Equation (4)*** under a wide range of possible values of the elastic parameters of the system (see Materials and methods). To further extend the scope of our equilibrium analytical model, in ***Appendix 1*** we discuss and analyze a fully dynamic, computational model of transport intermediate formation.

In order to compute the optimal membrane shape and TANGO1 capping state, we will seek for the minimum of the total free energy per unit area, ***Equation (4)***, as a function of the two degrees of freedom of the model: the capping fraction, *ω*, and the transport intermediate shape. For the latter, we use a dimensionless parameter to characterize the elongation state of the transport intermediate: *η = h*/2*R*, which is the height of the carrier divided by the diameter of a fully formed bud (***Figure 1–figure supplement 1B***). Taking this into account, we can write down the free energy per unit area, ***Equation (4)***, as (see Materials and methods):

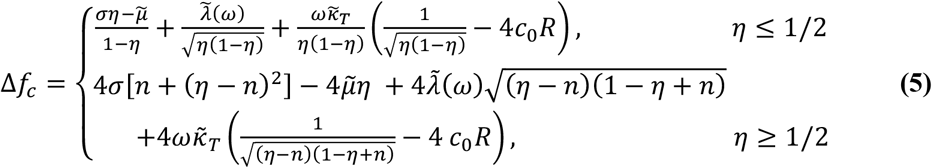

where 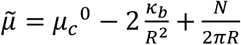 is the effective chemical potential, which depends on the binding energy of the coat to the membrane, *μ_c_*^0^, the bending rigidity of the membrane, *κ_b_*, and the applied pulling/pushing force, *N*; 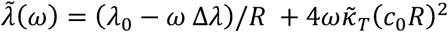, is the effective line tension of the coat; 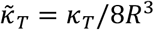 is the renormalized bending rigidity of the TANGO1 filament; and *n* is the number of fully-formed buds (see Materials and Methods). From the expression for the effective chemical potential, 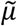, we can see that applying a force in the bud growth direction, *N*, plays the same role as increasing the coat binding free energy, *μ_c_*^0^, and therefore helps counterbalance the elastic resistance of the membrane to deformation.

The free energy per unit area of the transport intermediate, Δ*f_c_*, has a non-trivial dependence on the shape of the carrier, parametrized by the shape parameter, *η*, as given by ***Equation (5)***. This implies that multiple locally stable shapes, corresponding to different local minima of the free energy, can coexist. To illustrate this dependence, the profiles of the free energy per unit area, Δ*f_c_*, as a function of the shape parameter *η*, for both complete capping (TANGO1 rings forming around COPII patches), *ω* = 1, and no capping (no TANGO1 rings around COPII patches), *ω* = 0, are shown in ***Figure 2*** for two different scenarios (see also ***Figure 2-figure supplement 1*** for the detailed plot of the different contributions to the free energy of the system). In the first one, which corresponds to a situation where the COPII binding energy is relatively small, *μ_c_*^0^ = 0.03 *k_B_T/nm*^2^ (***Figure 2A***), the global minimum of the free energy corresponds to a shallow bud surrounded by a TANGO1 ring (depicted in green in ***Figure 2A***). Other locally stable shapes, corresponding to a shallow bud connected to a set of spheres, can be found (depicted in yellow in ***Figure 2A***). By contrast, in the second scenario illustrated in ***Figure 2B***, which corresponds to a situation of a relatively large COPII binding energy, *μ_c_*^0^ = 0.034 *k_B_T/nm*^2^, the transport intermediate will grow from an initially unstable shallow bud (depicted in red in ***Figure 2B***) to a locally stable, almost fully formed spherical carrier (depicted in yellow in ***Figure 2B***). Then, overcoming an energy barrier will result in further growth of the carrier into a large transport intermediate (depicted in green in ***Figure 2B***).

**Figure 2.**
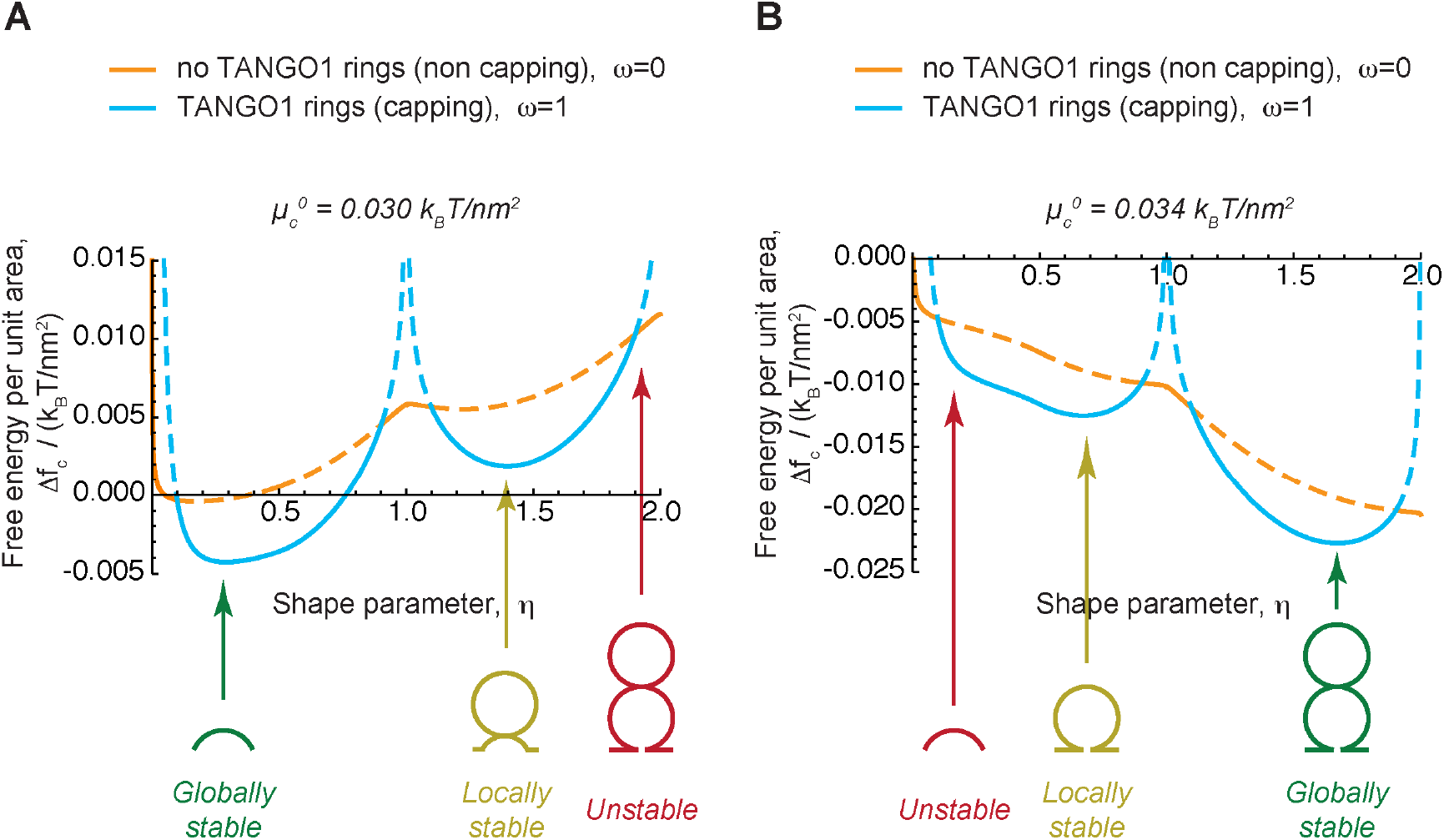
Free energy profile of a transport intermediate as a function of its shape and TANGO1 capping. **(A,B)** The free energy per unit area of the transport intermediate-TANGO1 system, Δ*f_c_*, plotted as a function of the shape parameter, *η*, for the COPII coat binding energy, 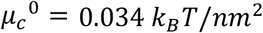 **(A)**, or 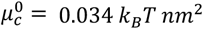 **(B)**. Two curves, corresponding to full COPII capping by TANGO1 (*ω* = 1, light blue curves) and full uncapping (*ω* = 0, orange curves), are shown in each panel. For each value of the shape parameter, *η*, the locally stable state of TANGO1 capping/unapping (lower free energy) is represented by the corresponding solid curve, whereas dashed curves indicate unstable states (higher free energy). A schematic representation of the shape of the transport intermediate for different values of the shape parameter, *η*, is depicted, including globally stable shapes (in green), locally stable (metastable) shapes (in dark yellow), as well as examples of unstable shapes (in red). We considered no applied force, *N* = 0, and the rest of the elastic parameters used for the calculations are specified in ***Table 1***.

### CAPPING-UNCAPPING TRANSITION OF TANGO1

Under which conditions do TANGO1 rings form by capping the edge of COPII patches? The free energy in ***Equation (4)*** has a linear dependence on the capping fraction, *ω*, and therefore is a monotonic function with respect to this variable. This implies that energy minimization will drive the system to either complete TANGO1 filament capping the edge of the COPII patch (*ω* = 1), or complete absence of TANGO1 around COPII (*ω* = 0), depending on the sign of *∂*Δ*f_c_/∂ω*. Partial rings, as experimentally observed (Raote *et al.*, 2018), might represent free TANGO1 filaments, rings in the process of assembly or disassembly, or complete rings complemented by other TANGO1-family proteins. The capping-uncapping transition corresponds, for a given value of the COPII boundary radius, *ρ*, to a stationary point of the free energy, ***Equation (4)***, with respect to the capping fraction, *∂Δf_c_/∂ω* = 0. This condition sets a critical value of the COPII boundary radius, *ρ*^*^, which defines the capping-uncapping transition,

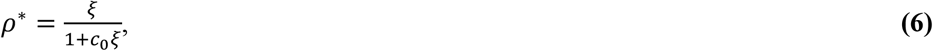

for 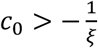, where 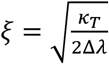 is a TANGO1-related length-scale, which, for the standard parameters described in ***Table 1*** is *ξ* ≈ 22 *nm*. For *ρ > ρ*^*^, there is complete capping of COPII domains by TANGO1 and thus full formation of TANGO1 rings; whereas for *ρ < ρ*^*^, there is complete absence of TANGO1 filaments around COPII domains and no TANGO1 rings are formed there (***Figure 3A***). Next, if we consider the case where the bud opening radius *ρ* is equal to the radius of curvature imposed by the spherical polymerization of COPII, *R*, we can define a critical value of the TANGO1 linactant strength below which there is no formation of COPII-capping TANGO1 rings, 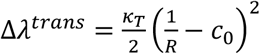 (***Figure 3A***).

**Figure 3.**
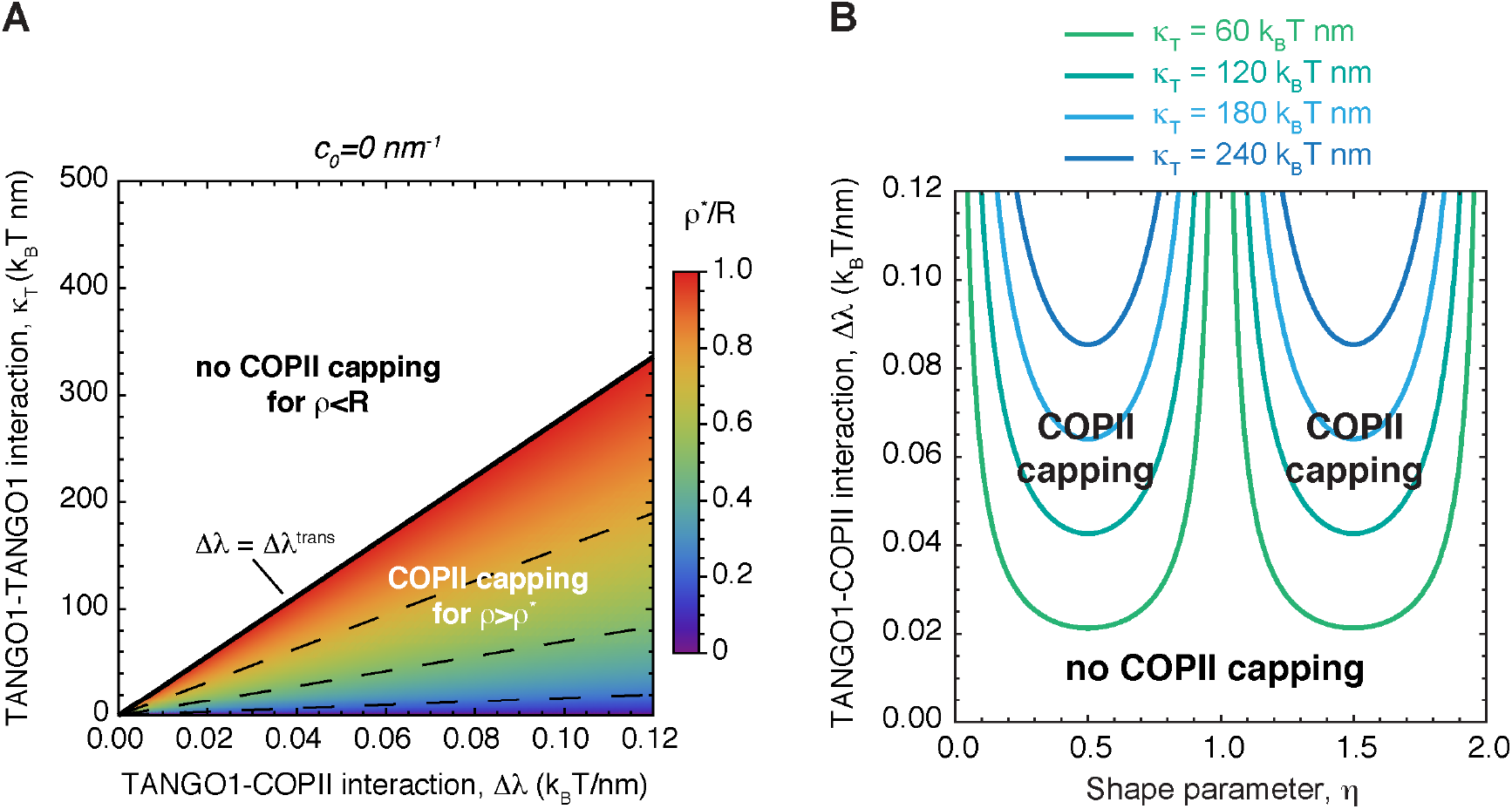
A capping-uncapping transition controls the formation of TANGO1 rings around COPII coats. **(A)** The capping-uncapping phase diagram as a function of the TANGO1 ring size, *ρ* (color-coded), given by ***Equation (6)***, is plotted against two TANGO1 filament parameters: the linactant strength of the TANGO1 filament, Δ*λ*, associated with the TANGO1-COPII interaction, and the filament bending rigidity, *κ_T_*, associated with the TANGO1-TANGO1 interaction, for a filament of zero spontaneous curvature.The critical value of TANGO1 linactant strength, Δ*λ^trans^*, below which no functional TANGO1 rings form (no COPII capping) is marked by the thick black line. **(B)** Capping-uncapping transitions of TANGO1 as a function of the shape of the transport carrier, *η*, plotted against the TANGO1 linactant strength, Δ*λ*, for different values of the filament bending rigidity, *κ_T_*.

However, whether a TANGO1 ring forms around a growing COPII patch depends not only on the TANGO1 properties (bending rigidity and linactant strength), but also on the actual shape of the COPII coat boundary (neck radius *ρ*) and therefore on the capacity of the coat to bend membranes, according to ***Equation 5***. Inspecting the free energy profiles shown in ***Figure 2*** we observe that during a quasi-static growth of the transport intermediate from *η* = 0 to the shape that corresponds to the global minimum of the system free energy (green shapes in ***Figure 2***), the system goes through a series of capping-uncapping transitions at *η = η^tr^*. These transitions occur when Δ*f_c_*(*η^tr^, ω* = 0) = Δ*f_c_*(*η^tr^, ω* = 1), which, from ***Equation (5)*** correspond to 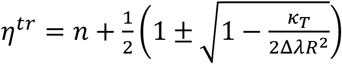, for *c*_0_ = 0 (***Figure 3B***). As expected, capping of COPII components by TANGO1 is promoted by large values of the TANGO1-COPII interaction, Δ*λ*, and prevented by large TANGO1 filament bending rigidities, *κ_T_* (***Figure 3B***). The general case for an arbitrary filament preferred curvature, *c_t_*, is analogous to the values of the opening angle described by ***Equation (6)***. From the free energy profile shown in ***Figure 2B***, we can also notice that the system needs to overcome an energy barrier to reach the globally stable state, so the system can be kinetically trapped into a locally stable deep bud (yellow shape in ***Figure 2B***). How such transitions occur is beyond the scope of this simple analytical model, since shape transition could proceed through transient uncapping of the TANGO1 filament or through intermediate shapes between a cylindrical tube and a set of spherical vesicles joined by a narrow connection, such as unduloids (see Materials and Methods). A more elaborate computational analysis of such intermediate shapes is presented in ***Appendix 1***. Taken together, our analysis shows that the growth of a transport intermediate can be modulated by the formation of TANGO1 rings capping COPII components, and a series of capping-uncapping transition should occur during the growth of a transport intermediate.

### FUNCTIONAL TANGO1 RINGS MODULATE TRANSPORT INTERMEDIATE FORMATION

To understand how TANGO1 rings modulate the formation of transport intermediates, we next sought locally and globally stable transport intermediate shapes. To this end, we computed the local minima of the overall energy of the system per unit area, ***Equation (5)***, for both single buds (shape parameter *η* < 1) or for large transport intermediates (shape parameter *η* > 1). In ***Figure 4***, we show the optimal shape of the intermediate –as measured by the optimal shape parameter, *η*^*^ − for a wide range of the model’s parameters. We first studied how transport intermediate formation depends on two key parameters: the COPII binding energy, *μ_c_*^0^, and the tension of the membrane, *σ* (***Figure 4A***). Our results show that both *(i)* increasing the ability of COPII to bind, and hence bend, membranes and *(ii)* decreasing the membrane tension favor the formation of large transport intermediates. Interestingly, for the range of COPII binding energies, 0.0285 *k_B_T/nm*^2^ < *μ_c_*^0^ < 0.0315 *k_B_T/nm*^2^, we notice that decreasing the tension of the ERES can lead to the elongation and of the transport intermediate from a shallow bud towards a pearled structure (***Figure 4A***, right panel), hence opening the possibility that membrane tension regulation at the procollagen export sites can induce the elongation of a transport intermediate. Interestingly, these results are in a good qualitative agreement with our dynamic model of transport carrier formation (see ***Appendix 1***). Next, we studied the role of a point-like force applied in the direction of bud growth. Our results show that the existence of such a force facilitates the transition from a shallow bud to a large intermediate (***Figure 4B,C***).

**Figure 4.**
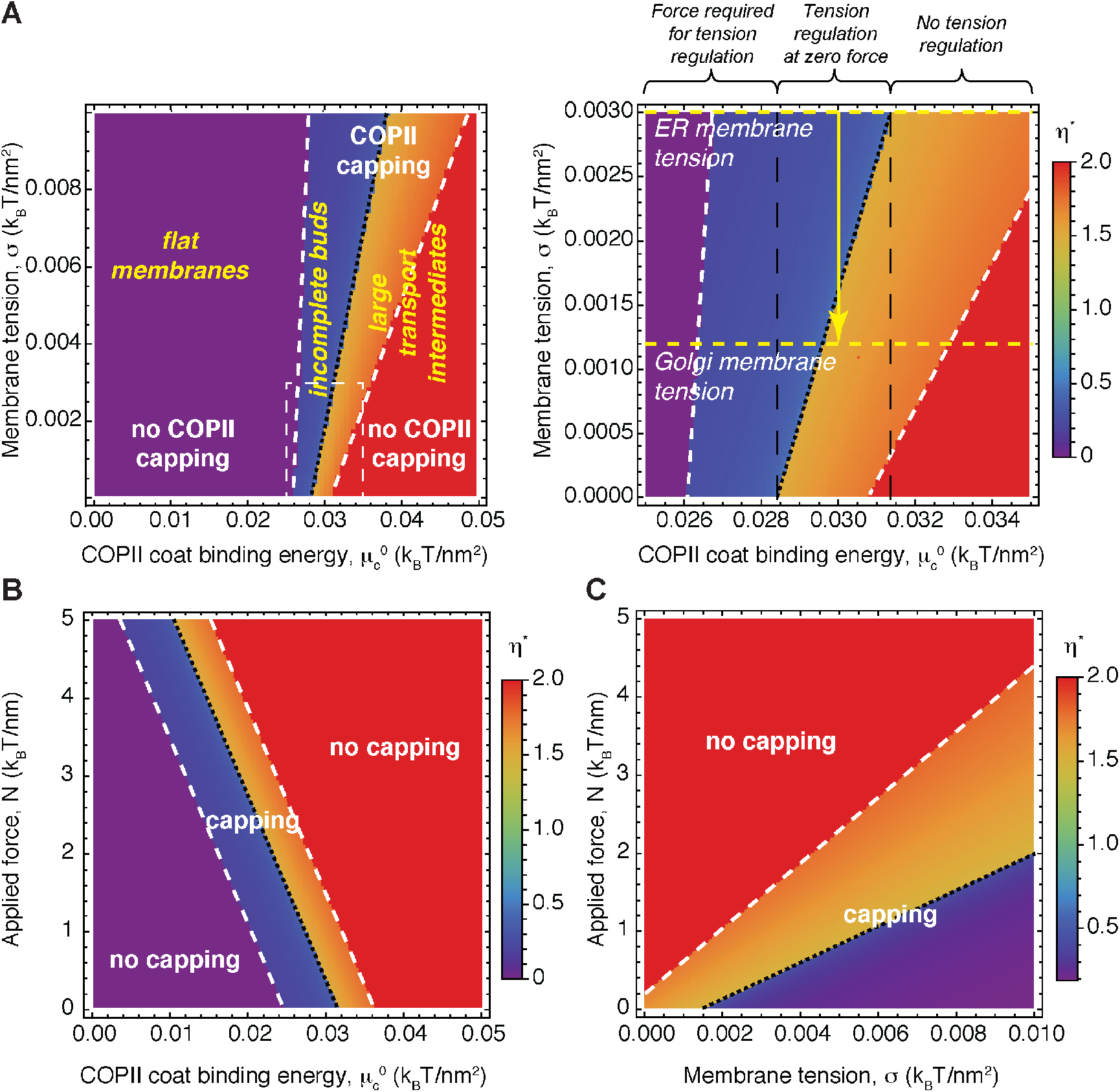
Shape diagram of the transport intermediate as a function of the membrane and COPII-controlled elastic parameters. **(A–C)** Two-dimensional shape diagrams indicating the shape of minimal elastic energy, represented by the optimal shape parameter, *η*^*^ (color-coded), and the state of TANGO1 capping (capping/uncapping transitions marked by thick, dashed, white lines). **(A)** Shape diagram plotted as a function of the COPII coat binding energy, 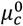, and of the membrane tension, *σ*, for vanishing applied force (*N* = 0). A zoom of the thin, dashed, white box on the left panel is shown in the right panel for clarity. Different shape transitions can be identified, from flat membranes, *η*^*^ = 0, to incomplete buds, *η*^*^ < 1/2, to large transport intermediates, *η*^*^ > 1, for both COPII capping (*ω* = 1) or not (*ω* = 0) by TANGO1. On the right panel, we mark the measured values for the standard ER and Golgi membrane tensions, and marked the regions of the shape diagram where tension regulation can or cannot describe the elongation of a shallow bud into a large transport intermediate (see text for details). **(B,C)** Shape diagram plotted as a function of the applied force, *N*, and the COPII coat binding energy, 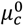, for *σ = σ_ER_* = 0.003 *k_B_T/nm*^2^ **(B)**; or the membrane tension, *σ*, for 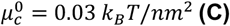. Unless otherwise specified, the elastic parameters used for all the calculations shown in **(A-C)** are listed in ***Table 1***.

Next, we computed how the properties of the TANGO1 filament, namely, the bending rigidity, *κ_T_*, and preferred curvature, *C*_0_, of the filament (given by the TANGO1-TANGO1 interactions), as well as the linactant strength (given by the TANGO1-COPII interactions), Δ*λ*, affect the formation of transport intermediates. Our results indicate that neither the linactant strength of the filament (***Figure 5A***) nor the rigidity (***Figure 5B***) or preferred curvature (***Figure 5-figure supplement 1***) of the TANGO 1 filament alter the transition zone between shallow buds (*η* < 1/2) and large intermediates (*η* > 1). However, filament properties are fundamental in controlling the capping/uncapping transition. Hence, in conditions of TANGO1 capping COPII components (such as large linactant strength or small filament bending rigidity), open shapes are favored as compared to flat or fully budded shapes (***Figure 5A,B*** and ***Figure 5-figure supplement 1***), suggesting once more that TANGO1 filaments can act as a means to prevent premature carrier fission before procollagen gets fully packaged into the nascent intermediate.

**Figure 5.**
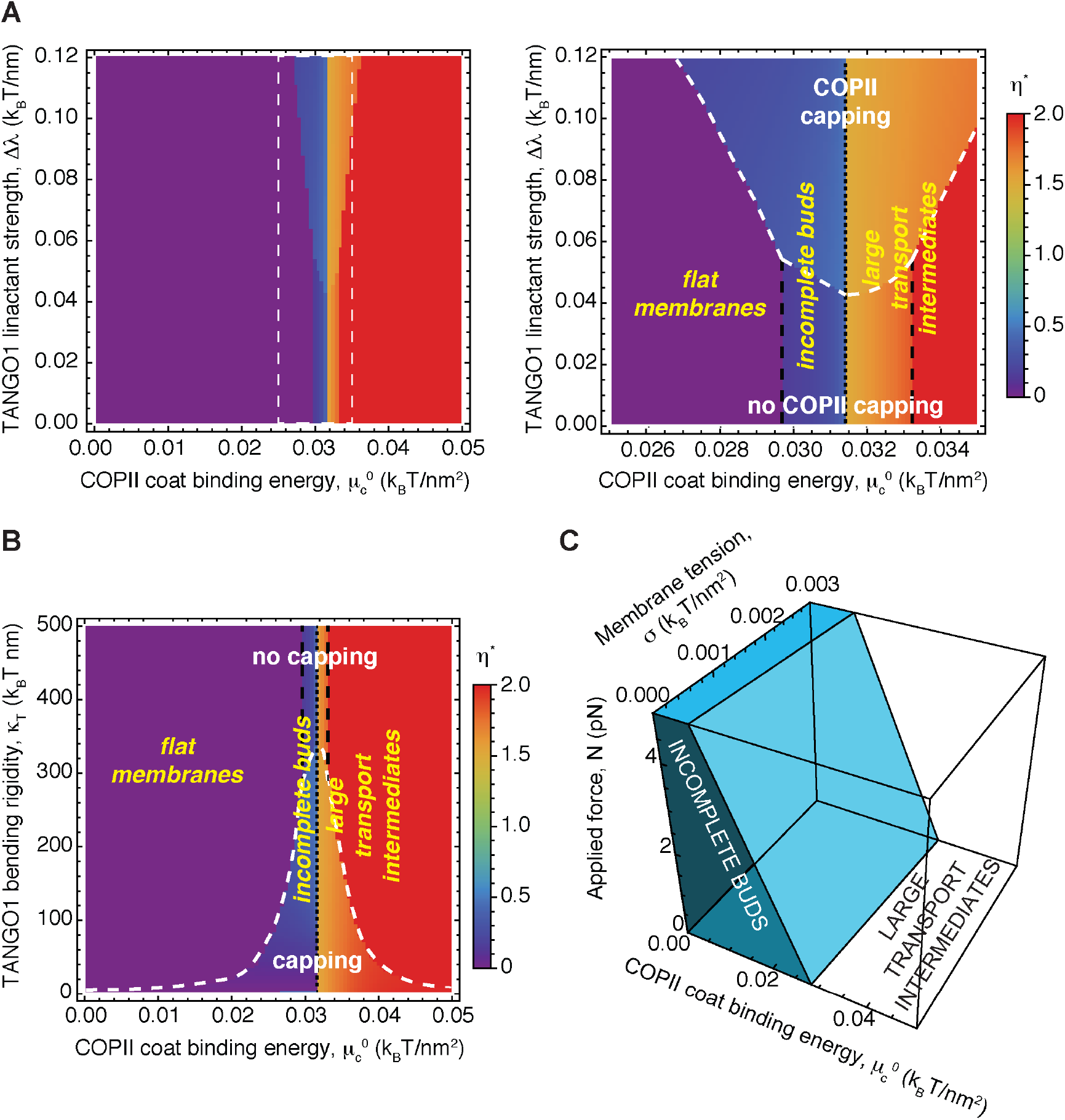
Shape diagram of the transport intermediate as a function of the TANGO1-regulated elastic parameters. **(A,B)** Two-dimensional shape diagrams indicating the shape of minimal elastic energy, represented by the optimal shape parameter, *η*^*^ (color-coded), and the state of TANGO1 capping (capping/uncapping transitions marked by thick, dashed, white lines). **(A)** Shape diagram plotted as a function of the COPII coat binding energy, 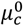, and of the TANGO1 linactant strength, Δ*λ*, for vanishing applied force (*N* = 0). A zoom of the thin, dashed, white box on the left panel is shown in the right panel for clarity, where the different shape transitions can be identified, from flat membranes, *η*^*^ = 0, to incomplete buds, *η*^*^ < 1/2, to large transport intermediates, *η*^*^ > 1, for both COPII capping (*ω* = 1) or not (*ω* = 0) by TANGO1. **(B)** Shape diagram plotted as a function of the COPII coat binding energy, 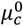, and of the TANGO1 filament bending rigidity, *κ_T_*, for vanishing applied force (*N* = 0). **(C)** Three-dimensional shape diagram, indicating the transition zone between incomplete buds (including both flat membranes, *η*^*^ = 0, and shallow buds, *η*^*^ < 1/2) and large transport intermediates (*η*^*^ > 1), as given by ***Equation (7)***, as a function of the COPII coat binding energy, 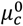; the membrane tension, *σ*; and the applied force, *N*. Unless otherwise specified, the elastic parameters used for all the calculations shown in **(A-C)** are listed in ***Table 1***.

Interestingly, based on ***Equation (5)***, we can have a good analytical estimate of the transition between incomplete shallow buds and large transport intermediates by considering the situation where the free energy of an incomplete bud with a certain shape (given by the height parameter *η)* equals to the free energy of a large intermediate with an extra pearl (given by the height parameter *η* + 1):

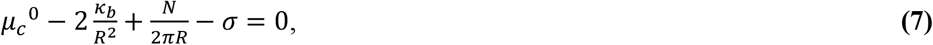

which is independent to the structural details of the TANGO1 filaments, and is shown in ***Figure 5C***. This expression allows us to define a critical force 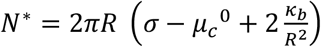; a critical coat binding energy, 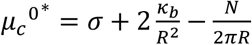; and a critical membrane tension, 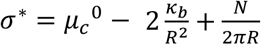; such that for 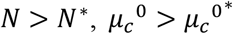, or *σ < σ*^*^, the transition to large transport intermediates is triggered. Taking the known or estimated parameters for the standard membrane tension of the ER, *σ_ER_* = 0.003 *k_B_T/nm*^2^ (Upadhyaya and Sheetz, 2004); for the membrane bending rigidity, *κ_b_* = 20 *k_B_T* (Niggemann, Kummrow and Helfrich, 1995); and for the size of the standard spherical COPII vesicle, *R* = 37.5 *nm* (Miller and Schekman, 2013) (see ***Table 1***); we get 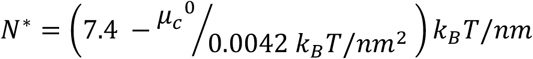 at *σ = σ_ER_*; *μ_c_*^0*^ = 0.031 *k_B_T/nm*^2^ at *σ = σ_ER_* and zero applied force (*N* = 0); and *σ*^*^ = *μ_c_*^0^ − 0.028 *k_B_T/nm*^2^, at zero force. Taken together, our results indicate that the formation of large transport intermediates can be favored by *(i)* an increase in the COPII coat binding energy, *μ_c_*^0^; *(ii)* a force, *N*, applied towards the direction of carrier growth; and/or a local decrease in the membrane tension, *σ*.

### MEMBRANE TENSION REGULATION BY TANGO1 CAN MEDIATE PROCOLLAGEN EXPORT

At present, we have no direct experimental evidence for the existence of a directed force pulling on growing buds at ERES, nor in the change of the COPII binding energy to assist in the formation of large transport intermediates. However, as mentioned earlier, TANGO1 rings serve to recruit COPI-coated membranes from the ERGIC (either vesicles or the ERGIC itself) to procollagen enriched export sites, which, upon fusion, could locally and transiently decrease the tension of the ERES. For this reason, we analyze here in more detail the requirements for the formation of large transport intermediates as a result of local reduction of the membrane tension from an initial value, *σ*_0_, to a lower value, *σ = σ*_0_ − Δ*σ*. Hence, ***Equation (7)*** can be written as 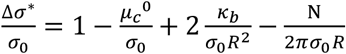. In ***Figure 6-figure supplement 1A,B***, we show the relative membrane tension reduction required to lead to the extrusion of long transport intermediates, 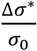, as a function of the COPII coat binding energy, *μ_c_*^0^, in the absence or presence of an applied force. The critical relative surface tension reduction defines three distinct regions in the 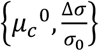 parameter space (***Figure 4A***, and ***Figure 6-figure supplement 1A***). At zero applied force, the first one corresponds to low values of COPII coat binding energy, *μ_c_*^0^ < 0.0284 *k_B_T/nm*^2^, where COPII polymerization energy is not large enough to allow growth of large transport intermediates, even in the absence of any lateral membrane tension (***Figure 4A***, left region in the right panel diagram, and ***Figure 6-figure supplement 1A***). Within the second region, which corresponds to intermediate values of the coat binding energy, 0.0284 *k_B_T/nm*^2^ < *μ_c_*^0^ < 0.0314 *k_B_T/nm*^2^, a partial reduction in membrane lateral tension can trigger formation of large transport intermediates in the absence of any applied force (***Figure 4A***, middle region in the diagram, and ***Figure 6-figure supplement 1A***). We refer to this intermediate region of the shape diagram as the region of transport intermediate formation triggered by TANGO1-mediated membrane tension regulation. Our calculations show that the relative decrease in membrane tension required to form large intermediates depends on the actual value of COPII coat binding energy (***Figure 6-figure supplement 1A***). We first consider the situation where tension in the ERES decreases from experimentally measured value of the ER membrane, *σ_ER_* = 0.003 *k_B_T/nm*^2^, to experimentally measured value of Golgi membranes, *σ_PS_* = 0.0012 *k_B_T/nm*^2^ (Upadhyaya and Sheetz, 2004), which accounts for a relative reduction in membrane tension of 60%. Such a reduction in tension can drive elongation of a bud to a large intermediate for values of the coat binding energy, *μ_c_*^0^ > 0.0296 *k_B_T/nm*^2^ (***Figure 6-figure supplement 1A***, red arrow). However, for more modest reductions in membrane tension, our results indicate that generation of large transport intermediates can also be triggered in the absence of a pulling force (***Figure 4A***, and ***Figure 6-figure supplement 1A***). Finally, in the third region of the shape diagram, which corresponds to large values of the coat binding energy, *μ_c_*^0^ > 0.0314 *k_B_T/nm*^2^, the polymerization of long COPII-coated tubular intermediates can occur spontaneously without the need of membrane tension reduction (***Figure 4A***, right region in the diagram, and and ***Figure 6-figure supplement 1A***). Remarkably, the region where membrane tension regulation can trigger the formation of large transport intermediates, corresponds to values of COPII binding energy that are of the same order of magnitude as the values measured for clathrin (*μ_clathrin_*^0^ = 0.024 ± 0.012 *k_B_T/nm*^2^) (Saleem *et al*., 2015). This suggests that our proposed mechanism of generating large intermediates by TANGO1-mediated membrane tension regulation is, at the very least, a physiologically feasible mechanism.

We next examined in more detail at the shape of the transport intermediate for these two different membrane tensions (*σ = σ_ER_* and *σ = σ_GC_*) as a function of the COPII coat binding energy within the region of membrane tension regulation (***Figure 6-figure supplement 1C***). As previously noted (***Figure 2***), multiple locally stable shapes exist, which are separated by free energy barriers (***Figure 6-figure supplement 1D***). Our computations indicate that a reduction in the membrane tension changes the free energy profile in such a way that globally stable shallow buds (solid lines, ***Figure 6-figure supplement 1C***) are converted into locally, but not globally, stable shapes (dashed lines, ***Figure 6-figure supplement 1C***). The free energy barrier for the transition to the new globally stable large transport intermediate, Δ*f*_1,2_, (***Figure 6-figure supplement 1D***) is decreased upon tension reduction (***Figure 6-figure supplement 1E***, black arrow), hence helping in the elongation of the transport intermediate.

Based on our previous experimental finding that functional TANGO1 rings recruit ERGIC53-containing membranes for procollagen export (Santos *et al*., 2015; Raote *et al*., 2018), we next aimed at understanding how membrane tension regulation can contribute in carrier elongation under physiological conditions. We computed the locally and globally stable shapes of the carrier, using our equilibrium model, for different tensions of the ER membrane for a defined value of the COPII coat binding energy within the tension regulation region (see ***Figure 4A***). Our results indicate that, for a wide range of membrane tensions, both shallow buds and large transport intermediates can be locally stable shapes (***Figure 6A***). For large values of the membrane tension, including the measured value for the ER membrane tension, *σ_ER_* = 0.003 *k_B_T/nm*^2^, bud growth is prevented and the globally stable shape corresponds to open, TANGO1-stabilized shallow buds (***Figure 6A***). For small values of the membrane tension, including the measured value of the Golgi membrane tension, *σ_GC_* = 0.0012 *k_B_T/nm*^2^, a large, elongated carrier is energetically favorable as compared to the shallow bud (***Figure 6A***). Importantly, transitions between the two locally stable shapes (shallow buds and large transport intermediates) are separated by free energy barriers, which, if too large, kinetically prevent shape transitions (***Figure 6-figure supplement 1D***). We then calculated the value of both free energy barriers: the one for the transition from a shallow bud to a large transport intermediate, Δ*f*_1,2_; and the other one for the opposite transition from a large intermediate to a shallow bud, Δ*f*_2,1_ (***Figure 6B*** and ***Figure 6-figure supplement 1D***). Our results indicate that reduction of the membrane tension parallels the reduction of the free energy barrier for carrier growth, as expected (***Figure 6B***). To estimate whether the free energy barrier reduction leads to a kinetically feasible transition to large transport intermediates at low membrane tension, we computed the overall free energy barrier in the projected area, Δ*F*_1,2_ = Δ*f*_1,2_*A_p_*, where *A_p_ = πR*^2^. For the conditions detailed in ***Figure 6***, we have a total free energy barrier for carrier elongation of Δ*F*_1,2_(*σ_ER_*) ≈ 33 *k_B_T* and Δ*F*_1,2_(*σ_GC_*) ≈ 8 *k_B_T*. Assuming that the transition follows Arrhenius kinetics, *τ = t*_0_*e*^Δ*F*_1,2_/*k_B_T*^, with a characteristic time scale *t*_0_~1 *ms* (Campelo *et al*., 2017), we get that the average transition time is reduced from *τ*(*σ_ER_*)~10^9^ *min* to *τ*(*σ_GC_*)~0.1 *min*. According to these estimations, membrane tension reduction can induce the transition from shallow buds to large transport intermediates. Interestingly, the possible shape recovery from an elongated carrier back to a shallow bud when the membrane tension is brought back to the initial large ER tension is Δ*F*_2,1_(*σ_ER_*) ≈ 25 *k_B_T*. This estimation suggests that the shrinkage of the carrier upon tension recovery would be kinetically prevented. Upon fusion of the ERGIC53-containing membranes to the ER exit sites, the tension of the membrane decreases, after which, tension is brought back to the initial value by tension diffusion (Shi *et al*., 2018). Under such cyclic membrane tension changes, from *σ_ER_* to *σ_GC_*, and then back to *σ_ER_*, the system undergoes a hysteretic cycle (***Figure 6A***). Starting at *σ_ER_*, the stable shape of the carrier is that of a shallow bud surrounded by a TANGO1 ring (point *(1)* in ***Figure 6A***). Reduction of the membrane tension to *σ_GC_* upon fusion of ERGIC membranes leads to reduction of the free energy barrier for carrier growth (***Figure 6B***) and spontaneous transition to a large transport carrier (point *(2)* in ***Figure 6A***). Finally, bringing the membrane tension back to *σ_ER_* does not parallel the shrinkage of the carrier back to a shallow bud, and the system is kinetically trapped in a large transport intermediate shape, even at high membrane tension (point *(3)* in ***Figure 6A***).

**Figure 6.**
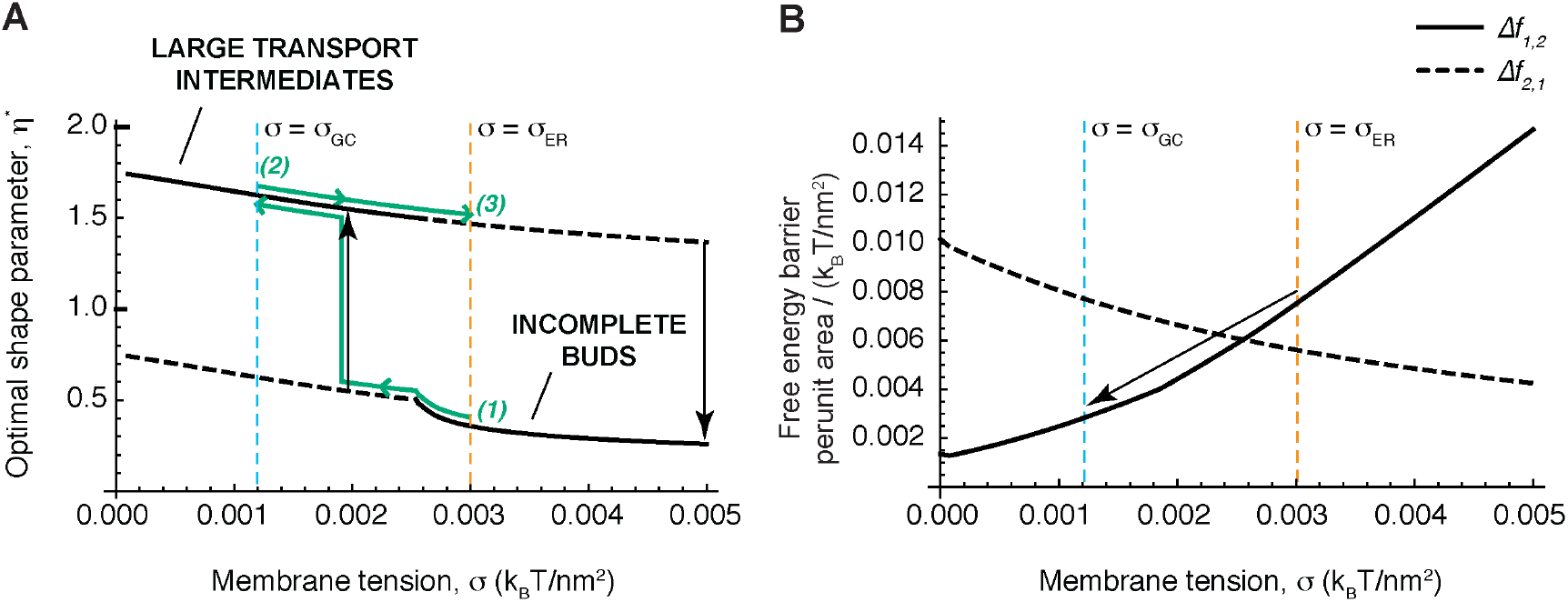
Membrane tension regulation by TANGO1 can mediate procollagen export. **(A)** The optimal shape parameter, *η*^*^, is plotted as a function of the membrane tension, *σ*, for a COPII coat binding energy, 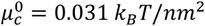, within the tension regulation regime (see ***Figure 4A*** and ***Figure 6–figure supplement 1A***). Globally stable shapes, for each case, are indicated by solid lines, whereas dashed lines represent locally, metastable shapes. A transient reduction in the membrane tension (green arrows from point (1) to point (2)) can lead to the growth of the transport intermediate (marked by the black vertical arrow, whereas recovery of the tension to the initial value (green arrows from point (2) to point (3)) can keep the system in a kinetically arrested metastable configuration. **(B)** The free energy barriers separating the incomplete bud from the large intermediate morphologies, Δ*f*_1,2_ (solid lines), and Δ*f*_2,1_ (dashed lines) (see ***Figure 6–figure supplement 1D*** for definitions), are plotted as a function of the membrane tension, *σ*, for the same parameters as used in **(A)**. The arrow illustrates how decreasing the membrane tension leads to a reduction of the energy barrier for growth of the transport intermediate. Unless otherwise specified, the elastic parameters used for all the calculations shown in **(A,B)** are listed in ***Table 1***. The values of the ER and Golgi membrane tensions are marked by dashed orange and blue lines, respectively.

### DYNAMIC COMPUTATIONAL MODEL OF TANGO1-ASSISTED TRANSPORT INTERMEDIATE FORMATION

The equilibrium model presented here (***Equation (4)***) allowed us to gain valuable insight into the physical mechanisms by which TANGO1 rings can deform the ER membrane for procollagen export. However, as any other model, ours is based in a number of assumptions and has its limitations. First, it is an equilibrium model, so the shapes of the carriers are found as those that locally minimize the free energy of the system. Hence, the dynamic behavior of shape transitions and protein redistribution cannot be fully grasped in a quantitative manner. Second, in our model we assumed that COPII coats polymerize as spherical lattices of fixed curvature. Indeed COPII coats usually adopt such shapes (Faini *et al*., 2013), but it has been recently reported that COPII coats can also adopt cylindrical arrangements (Zanetti *et al*., 2013; Ma and Goldberg, 2016). To overcome these two main limitations, we derived a dynamic model of TANGO1-assisted transport carrier formation, where there is no imposed constraint on the shape of the transport intermediate, besides that of imposing axisymmetric shapes (see ***Appendix 1*** for the detailed description and analysis of the dynamic model) (Tozzi, Walani and Arroyo, 2019). Importantly, this dynamic model qualitatively recapitulates the formation of TANGO1 rings and their ability to stabilize open carriers (see ***Appendix 1***), in accordance with our equilibrium model (***Figure 4***).

We exploited our dynamic model to understand whether TANGO1-mediated tension regulation can dynamically enable carrier elongation for procollagen export. To test this, we simulated the sequential decrease of membrane tension representative of the recruitment and fusion of ERGIC53-containing vesicles to the procollagen export site. To mimic the presence of procollagen molecules inside the growing carrier preventing the total closure of the neck (***Figure 1D-H***), we set a minimum neck radius threshold of 7.5 nm (see ***Appendix 1*** for details). Starting from a stable shallow bud of shape parameter, *η* − 0.6, at a large membrane tension of *σ* = 0.008 *k_B_T/nm*^2^ (***Figure 7D** (i)*), we apply a transient tension reduction as follows. First, the tension is reduced to *σ* = 0.004 *k_B_T/nm*^2^, leading to bud re-growth. Second, when the bud height reaches *η* = 1.5 (just after the start of neck closure), the membrane tension is gradually set back to its original value of *σ* = 0.008 *k_B_T/nm*^2^ over a 10 ms time ramp (***Figure 7A-C***). We find that the carrier reaches a new equilibrium state at *η* − 2.4, corresponding to a “key-hole” shape of about 170 nm height (***Figure 7D** (ii)*). Next, continuing from this new equilibrium state, we repeat the transient tension reduction protocol, this time initiating the ramp to recover high tension for *η* = 3.5, just after a new neck is formed. Once again, the transport intermediate evolves towards a new equilibrium “unduloid-like” shape at *η* − 4.2 (***Figure 7D** (iii)*, and ***Movie 1***). Interestingly, this corresponds to a height of about 300 nm, the typical length of a folded procollagen molecule.

**Figure 7.**
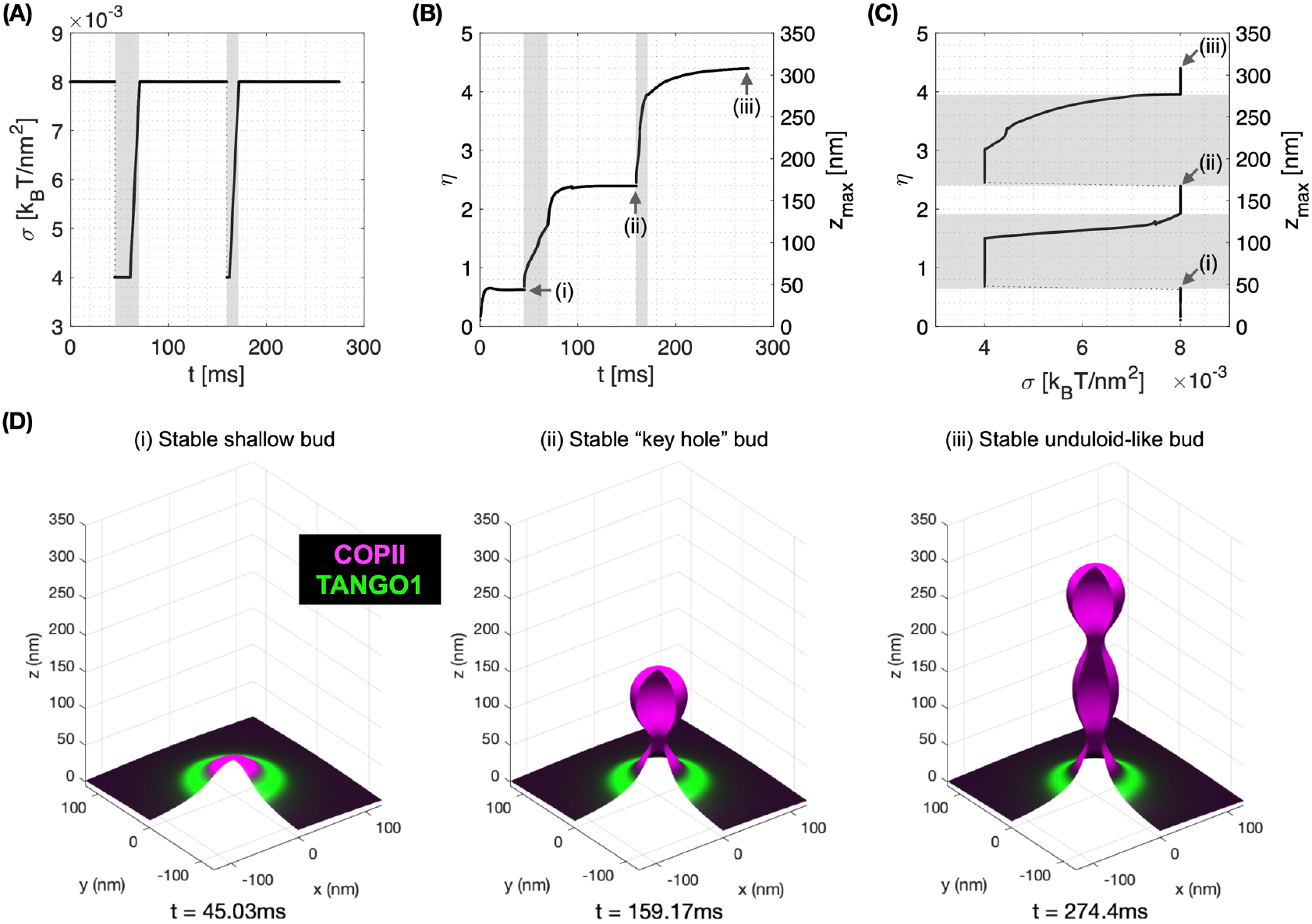
Membrane tension regulation by TANGO1 enables the formation of large procollagen-containing transport intermediates. A transient decrease of membrane tension mimicking ERGIC membrane recruitment by TANGO1 enables the sequential growth of COPII-TANGO1 transport intermediates. **(A)** Computational protocol for the applied membrane tension as a function of time. When an equilibrium state is reached, *σ* is transiently reduced from 0.008 to 0.004 *k_B_T/nm*^2^ until the transport intermediate grows to either *η* = 1.5 or *η* = 3.5. Then *σ* is progressively restored to 0.008 *k_B_T/nm*^2^ following a linear ramp of ~10 ms. **(B)** Resulting shape factor as a function of time. **(C)** Shape factor as a function of membrane tension. Gray regions in **(A-C)** correspond to low-tension regimes. **(D)** Three-dimensional rendering of the bud shape and protein surface distribution at the three equilibrium shapes indicated in panels **(B)** and **(C)**. Parameters as in ***Appendix 1–Table 1***, with *χ_c_* = −2 *k_B_T* (see ***Appendix 1*** for details, and corresponding ***Movie 1*** for a dynamic representation of the simulation).

Altogether, our results support the scenario by which TANGO1 self-organizes around COPII coats and stabilizes shallow buds at physiological membrane tensions. This mechanism is enabled by TANGO1 affinity to COPII, TANGO1 ring bending rigidity, and the modulation of COPII coat line energy. The stability of shallow buds might facilitate procollagen packaging as well as the recruitment of ERGIC membranes to the ERES. Transient membrane tension reduction, possibly mediated by fusing ERGIC membranes to the ER, allows the buds to grow from shallow to elongated pearled transport intermediates of sizes compatible with the encapsulation of procollagen molecules.

## DISCUSSION

### COMPARISON WITH EXPERIMENTAL RESULTS

We previously reported that the appearance of well-formed, regular TANGO1 rings at the ERES correlated with the ability of those cells to efficiently export procollagen from the ER (Raote *et al*., 2017, 2018). First, when the association of TANGO1 with COPII subunits was abrogated –either by expressing TANGO1-ΔPRD or by silencing the expression of SEC23A–, TANGO1 formed either smaller rings or long linear filamentous structures or planar clusters. Second, when the mutual association between individual TANGO1 filament components was hampered –either by expressing TANGO1-ΔCC2 or by silencing the expression of cTAGE5–, TANGO1 formed rings that were of less defined sizes and with more irregular, less circular shapes. And third, when the association of TANGO1 with the tethering factors that recruit ERGIC53-containing membranes to the ERES was inhibited–either by expressing TANGO1-ΔCC1, TANGO1-ΔTEER, or by silencing the expression of the tethering factors RINT1, NBAS, or ZW10–, TANGO1 rings were virtually absent at the ERES.

In cells expressing TANGO1-ΔPRD, the interaction between one of the filament components, TANGO1, and the COPII subunits is abolished, indicating that, although a TANGO1 filament could still be formed –this mutant does not alter the interaction between TANGO1 and other TANGO1 or cTAGE5 proteins (Raote *et al*., 2018)–, the filament should be less able to cap COPII components, and therefore less line-active, because the affinity to bind to the peripheral COPII subunits is reduced. In this situation the filament proteins cTAGE5 (Saito *et al*., 2011, 2014) and TANGO1-Short (Maeda, Saito and Katada, 2016) can still bind Sec23A and therefore reduce, albeit to a lesser extent than in wild-type cells, the COPII patch line energy. Our theoretical results presented here predict that decreasing the TANGO1 filament linactant strength, Δ*λ*, results in a lower chance of having open carriers (***Figure 5A***), which can lead to the observed defects in terms of ring structure and procollagen export.

In summary, the results we obtained from our physical model of large transport intermediate formation reinforce the notion that TANGO1 rings serve to control the formation of COPII carriers. TANGO1 rings can stabilize the COPII bud neck and thus prevent their premature closure by kinetically arresting or slowing down the completion of a spherical carrier. In such a situation, carrier expansion –according to the results of our model– can proceed via three different scenarios (***Figure 4***): *(i)* increase in the polymerization ability of COPII coats; *(ii)* appearance of a directed force applied at the growing carrier and pointing towards the cytosol; and *(iii)* local reduction of the membrane tension. TANGO1 can directly or indirectly control each of these possibilities (Ma and Goldberg, 2016; Raote *et al*., 2018). Interestingly, the TANGO1 ring properties, such as the linactant power of TANGO1 or the TANGO1 filament bending rigidity, are not drivers of the incomplete bud to large transport intermediate transition (***Figure 5***), but they seem to act more as kinetic controllers of the transition by preventing bud closure and stabilize open carrier shapes (***Figures 4,5***).

### PROPOSAL OF EXPERIMENTAL APPROACHES TO TEST THE MODEL

In this article, we proposed and analyzed a theoretical model to understand how TANGO1 molecules assemble into functional rings at the ERES, and how these rings can control the shape of transport intermediates. Our theoretical results have the potential to open up new avenues for experimental research on this topic and provide a common framework within which data and results can be understood. In particular, we envision that our work will stimulate future experimental efforts to test mechanisms of ERES organization and collagen export. We propose here some possible routes by which the hypotheses and predictions of our model as well as some of the open questions it raised could be experimentally tested.

#### Does TANGO1 form a linear or quasi-linear filament held together by lateral protein-protein interactions?

A first step to address this question will be to resolve the stoichiometry of the TANGO1 family proteins within a TANGO1 ring. Controlled photobleaching (Lee *et al*., 2012) or DNA-PAINT (Stein *et al*., 2019) of the single-labeled, endogenously-expressed proteins would allow the recording of the number and spatial positions of single fluorophores in individual TANGO1 rings. These results, after complete quantitative reconstruction of all the single molecule signals, should provide an absolute stoichiometry and ultra-resolved structure of TANGO1 organization in the ERES. Ultimately, *in vitro* reconstitution of TANGO1 ring formation in synthetic lipid bilayers by using recombinant proteins will be of paramount importance to experimentally observe the formation of TANGO1 filaments, assess the minimal components required for their formation, and eventually measure the elastic properties of a TANGO1 filament.

#### Is tension homeostasis, controlled by TANGO1-directed fusion of incoming ERGIC membranes, a mechanism for transport intermediate formation?

Future efforts in applying cutting-edge, super-resolution multicolor live-cell microscopy (Bottanelli *et al*., 2016; Ito, Uemura and Nakano, 2018; Liu *et al*., 2018; Schroeder *et al*., 2019) will help monitor the fusion of ERGIC membranes to the ER and couple these events to the formation of procollagen-containing transport intermediates. In addition, our hypothesis of TANGO1-mediated regulation of membrane tension is based upon the premise that fusion of ERGIC-53-containing vesicles to the procollagen export sites locally decreases the membrane tension. A recently-established fluorescent membrane tension sensor (Colom *et al*., 2018; Goujon *et al*., 2019) could provide a means to monitor such effects in relation to procollagen export.

#### Is there an outwards-directed force driving transport intermediate elongation?

It has been shown that procollagen export from the ER does not require the presence of an intact microtubule network (McCaughey *et al*., 2019), however the involvement of other force-producing agents, such as actin-myosin networks, remains unknown. The identification of physiologically meaningful interactors of TANGO1 by proximity-dependent labeling assays, such as BioID (Roux *et al*., 2018), and the subsequent screening for candidates that can exert those forces would set the grounds to identify possible molecular players involved in force-generation. However, it is important to stress that our model can explain formation of large transport intermediates even in the absence of an applied force (***Figure 6***).

#### Finally, what is the shape of the transport intermediate that shuttles collagens from the ER to the ERGIC/Golgi complex?

To this end, three-dimensional, multicolor super-resolution microscopy techniques, such as 3D single molecule localization microscopy (3D-SMLM) or 3D stimulated emission depletion (3D-STED) microscopy, could provide sufficient resolution to map the three-dimensional morphology of the transport intermediates. Recent efforts by using 3D-SMLM and correlative light and electron microscopy (CLEM) have revealed the existence of large procollagen-containing structures (Gorur *et al*., 2017; Yuan *et al*., 2018). However, a recent report suggested that those structures were directed for lysosomal degradation and not for trafficking to the Golgi complex (Omari *et al*., 2018). By contrast, direct transport of procollagen between the ER and the Golgi complex by a short-loop pathway in the absence of large vesicles has been recently proposed (McCaughey *et al*., 2019), opening to the possibility of a direct tunneling mechanism for trafficking proteins between compartments (Raote and Malhotra, 2019). Eventually, the use of modern electron microscopy techniques such as cryo-electron tomography (Beck and Baumeister, 2016) or focused ion beam-scanning electron microscopy (FIB-SEM) (Nixon-Abell *et al*., 2016) will help solve this issue on the morphology of the transport intermediates that shuttle procollagens form the ER to the Golgi complex.

### TANGO1 AS A REGULATOR OF MEMBRANE TENSION HOMEOSTASIS

We previously showed that TANGO1 forms circular ring-like structures at ERES surrounding COPII components (Raote *et al*., 2017). We also revealed interactions that are required for TANGO1 ring formation, which are also important to control TANGO1-mediated procollagen export from the ER (Raote *et al*., 2018). However, it still remained unclear how TANGO1 rings organize and coordinate the budding machinery for efficient procollagen-export. Here, we proposed, described, and analyzed a feasible biophysical mechanism of how TANGO1 mediates the formation of procollagen-containing transport intermediates at the ER. The results of our model suggest that TANGO1 rings serve as stabilizers of open buds, preventing the premature formation of standard COPII coats. TANGO1 is ubiquitously expressed in mammalian cells, including cells that secrete very low amounts of collagen. Furthermore, TANGO1 resides in most ERES in all these different cell lines, yet small COPII-coated carriers should form normally in those sites. How can this be understood? We propose that the ability of TANGO1 to form a ring at COPII-coated ERES is a first requirement for TANGO1 to promote procollagen export in non-standard COPII-dependent transport intermediates. Accumulations of export-competent procollagen at the ERES could re-organize the TANGO1 molecules laying there into functional rings surrounding COPII components and kinetically preventing the formation of small COPII carriers. Tethering of ERGIC53-containing vesicles mediated by the TANGO1 TEER domain (Raote *et al*., 2018) could be the trigger to allow for carrier growth. Importantly, the ER-specific SNARE protein Syntaxin18 and the SNARE regulator SLY1, which together trigger membrane fusion at the ER, are also required for procollagen export in a TANGO1-dependent manner (Nogueira *et al*., 2014). Fusion of ERGIC membranes to the sites of procollagen export would lead to a local and transient reduction of the membrane tension (Sens and Turner, 2006), which can promote, according to our theoretical results, the growth of the COPII carrier. We proposed that ERGIC membranes fuse directly to the growing transport intermediate to allow for membrane addition and tension release; and showed that compartment mixing can be arrested by the TANGO1 ring serving as a diffusion barrier (Raote *et al*., 2020). In this scenario, TANGO1 would act as a regulator of membrane tension homeostasis to control procollagen export at the ERES. In parallel, we can also hypothesize a situation where TANGO1 rings help pushing procollagen molecules into the growing carrier and couple this pushing force to procollagen folding, through the chaperone HSP47 (***Figure 1F***). Because HSP47 chaperone assists in folding (and hence in rigidifying) procollagen, the physical interaction between TANGO1 rings and procollagen/HSP47 could serve as a means to couple procollagen folding to force production. Although the existence of this pushing force is largely speculative, it could, according to our model, promote formation of a large intermediate and hence TANGO1 could act as a sensor of procollagen folding to couple it with the export machinery.

Physically, our model can be understood in terms of a competition between different driving forces. Each of these forces can either prevent or promote the elongation of procollagen-containing transport intermediates. First, COPII coats normally polymerize following a spherical arrangement, and the forming COPII lattice has a propensity to reduce the amount of free edges. Binding of the TANGO1 PRD to these edge-localized peripheral COPII subunits has a two-fold effect: *(i)* it prevents binding of other COPII subunits that would complete the polymerization of the coat into a spherical vesicle, and thereby *inhibits* the fission of the vesicle; and *(ii)* by stabilizing the neck of an open carrier, TANGO1 forms a ring around COPII subunits. Eventually, the inhibition of premature bud fission and therefore the stabilization of open carriers converts a small bud into a large transport intermediate, and finally into a tunnel connecting ER to the ERGIC/early Golgi cisternae (Raote and Malhotra, 2019). In physical terms, the role of TANGO1-PRD in this process is to effectively reduce *line tension* of COPII coat (by acting as a linactant), without directly changing the *surface tension* of the underlying membrane. In our view, TANGO1-PRD is important to keep the transport intermediate connected to the ER by preventing fission of the nascent carrier. Subsequently, fusion of ERGIC membranes, which locally and transiently decreases the membrane surface tension, will lead, as our model predicts, to growth of a large transport intermediate (***Figure 1G,H***).

What controls the organelle size in the context of intracellular trafficking? There has been a lot of work on what set the size of organisms, the size of tissues in an organism, and the size of cells in a tissue. However there has been less work on the question of what sets the size of organelles relative to the cell. Extensive cargo transfer while trafficking bulky cargoes such as collagens leads to large amounts of membrane being transferred from one organelle to another. To maintain organellar homeostasis, loss of membrane from a compartment has to be concomitantly compensated by membrane acquisition from the biosynthetic pathway or by trafficking from other organelles; the arrival and departure of membrane at each compartment has to be efficiently balanced. How is this homeostatic balance controlled? Changes in membrane tension have been described to affect rates of exocytosis and endocytosis at the plasma membrane (Apodaca, 2002; Kosmalska *et al*., 2015; Wu *et al*., 2017). Interestingly, a theoretical model has also established a crucial role for membrane tension in modulation the transition to bud clathrin-coated vesicles (Hassinger *et al*., 2017). Furthermore, it has been recently proposed that Atlastin-mediated maintenance of ER membrane tension is required for the efficient mobility of cargo proteins (Niu *et al*., 2019). However, control of endomembrane trafficking by membrane tension is more challenging to study experimentally and hence still remains poorly understood. We propose that TANGO1 serves as a hub in the ER to connect different organelles by controlling the local tension homeostasis at specific membrane sub-domains and regulating the membrane flux between these organelles.

In summary, we proposed a theoretical mechanical model that explains how TANGO1 molecules form functional rings at ERES, and how these TANGO1 rings assemble the machinery required to form a large transport intermediate commensurate with the size of procollagens. We envision that our hypotheses and the predictions of our model will guide new lines of experimental research to delineate mechanisms of COPII coats organization for the export of complex cargoes out of the ER.

## MATERIALS AND METHODS

### DETAILED DESCRIPTION OF THE PHYSICAL MODEL OF TANGO1-DEPENDENT TRANSPORT INTERMEDIATE FORMATION

Here we present the detailed description and derivation of the physical model of TANGO1-dependent transport intermediate formation presented in the main text. Our model builds on a previously presented mechanical model for clathrin-coated vesicle formation (Saleem *et al*., 2015), which we extended to allow for the growth of larger transport intermediates by incorporating *(i)* the effects of TANGO1 rings on COPII coats; *(ii)* the reduction of the membrane tension by the tethering and fusion of ERGIC53-containing membranes; and *(iii)* an outward-directed force (***Figure 1–figure supplement 1A***).

Analogously to the clathrin vesicle model by Saleem et al. (Saleem *et al*., 2015), we consider that the free energy per unit area of coat polymerization onto the membrane, *μ_c_*, has a bipartite contribution arising from the positive chemical potential of COPII binding to the membrane, *μ_c_*^0^, and from the negative contribution of membrane deformation by bending, so 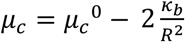, where *κ_b_* is the bending rigidity of the lipid bilayer, and *R* is the radius of curvature imposed by the spherically polymerized COPII coat. An additional term associated to the possible elastic deformation of the COPII coat could be considered as 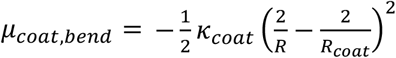, where *κ_coat_* is the coat rigidity and *R_coat_* is the spontaneous radius of curvature of the coat (Iglič, Slivnik and Kralj-Iglič, 2007; Boucrot *et al*., 2012). However, we assume that the coat is considerably more rigid than the membrane, *κ_coat_ ≫ κ_b_*, so there is no coat deformation and *R = R_coat_*. Alternatively, coat contribution to membrane bending has also been tackled by using a spontaneous curvature-based model (Agrawal and Steigmann, 2009; Hassinger *et al*., 2017). In our analytical model we follow the approach of Saleem et al. (Saleem *et al*., 2015), which allows us to define the preferred spherical architecture of the polymerized coat. A spontaneous curvature-based approach was followed for our computational analysis of carrier shapes (***Appendix 1***). In summary, the free energy per unit area of the initially undeformed membrane due to COPII polymerization, *f_coat_*, can be expressed as

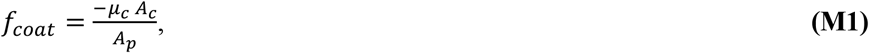

where *A_c_* is the surface area of the membrane covered by the COPII coat, and *AD* is the projected area of the carrier, that is, the area of the initially undeformed membrane under the carrier (Saleem *et al*., 2015) (***Figure 1–figure supplement 1B***). We also need to consider a line energy for the coat subunits laying at the edge of the polymerizing structure. This line energy per unit area reads as

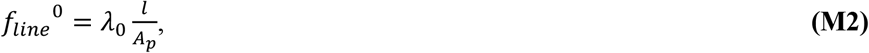

where *λ*_0_ is the line tension of the bare coat, and *l* = 2*πρ* is the length of the carrier edge, associated to the opening radius at the base of the carrier, *ρ* (***Figure 1–figure supplement 1B***). Next, we consider the contribution of the membrane tension, *σ*, to the free energy per unit area of the system, which reads as

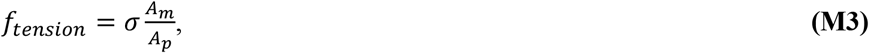

where *A_m_* is the surface area of the entire membrane after deformation.

The following step is to expand on this model to include the contributions by which TANGO1 can modulate the formation of a transport intermediate. TANGO1 filaments are described by their length, *L_T_*, and by their persistence length, 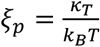, –where *κ_T_* is the filament bending rigidity and *k_B_T* is the thermal energy, equal to the Boltzmann constant times the absolute temperature (Doi and Edwards, 1986)–, which describes how stiff the filament is. As long as the filament length is not much larger than the persistence length, the bending energy of the TANGO1 filament can be expressed as 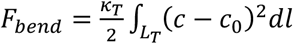, where *c* and *c*_0_ are the actual and spontaneous curvature of the filament, respectively, and the integral is performed over the entire filament length. We define positive spontaneous curvatures of the filament as those where the TANGO1-COPII interacting domains lie on the concave side of the filament, and negative when they lie on the convex side. For a TANGO1 filament of length *L_T_*, that is bound to the circular boundary length of a COPII patch (of radius *ρ*), the filament bending energy per unit length can be written as 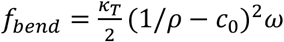, where we assumed that any existing filaments not adsorbed to the COPII patches adopt the preferred curvature, *c*_0_, and where *ω* is the capping fraction: the fraction of COPII domain boundary length covered (“capped”) by TANGO1 molecules. Hence, analogously to our discussion for the free energy of coat binding to the membrane, ***Equation (M1)***, we can write the free energy per unit area of a TANGO1 filament as

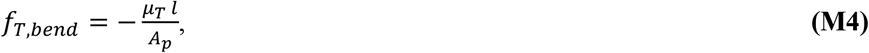

where *μ_T_ = −f_bend_* includes the negative contribution of the filament bending energy. A positive contribution of the filament assembly chemical potential, *μ_T_*^0^, is not considered here since we assume that the assembly chemical potential is independent of whether the filament is capping or not a COPII patch and hence the fraction of TANGO1 monomers forming a filament is independent of the capping fraction. Moreover, we want to stress that the bending energy penalty of the filament diverges when the bud approaches closure, meaning that either there is *uncapping* of the TANGO1 filament from the edge of the COPII coat at narrow necks or the shape transition of the carrier goes through intermediate shapes with a relatively large bud neck, such as Delaunay shapes (e.g. unduloids) (Naito and Ou-Yang, 1997). This second option is analyzed in ***Appendix 1***. In addition, TANGO1 proteins have an affinity to bind COPII components, and hence adsorb to the boundary of the COPII domains by binding the most external subunits. We therefore consider an extra free energy term associated to this TANGO1-COPII interaction, which is proportional to the boundary length of the COPII domain capped by TANGO1, and hence reads as

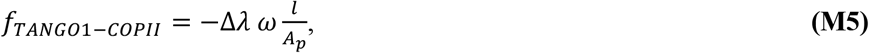

where Δ*λ* is the interaction strength between TANGO1 and COPII. We can write together ***Equations (M2)*** and ***(M5)*** as

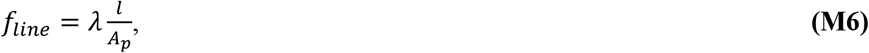

where 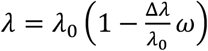 can be understood as the effective line tension of the COPII coat, in which Δ*λ/λ*_0_ is the relative reduction in the line tension due to TANGO1 capping, and hence is a measure of the linactant power of TANGO1.

Finally, the mechanical work performed by the outward-directed force, *N*, is also included in the free energy of the system, as

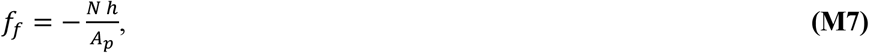

where *h* is the length of the carrier (***Figure 1–figure supplement 1B***). At this stage, for the sake of simplicity, we disregard the effects of the growth-shrinkage dynamics of the polymerizing COPII lattice. Hence, the total free energy per unit area of the carrier, *f_c_*, is the sum of all these contributions ***Equations (M1,3,4,6,7)***,

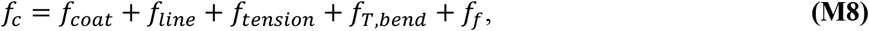

which is presented in ***Equation (4)*** in the main text.

#### Geometry of the problem

Based on the proposed geometries for the growing carrier we can distinguish three geometries, depending on how complete the transport intermediate is: shallow buds, deep buds, and pearled intermediates (***Figure 1–figure supplement 1B***, panels *(i)* to *(iii)*, respectively). These shapes will allow us to calculate as a function of the carrier morphology the geometric parameters that enter in ***Equation (4)***, namely, the area of the coat, *A_c_*, the area of the membrane, *A_m_*, the projected area, *A_p_*, and the opening radius at the coat rim, *ρ* (Saleem *et al.*, 2015). A convenient quantity to parametrize the shape of the carrier is the height of the carrier, *h*, which we will use in a dimensionless manner by normalizing it to the diameter of the spherical COPII bud, *η = h*/2*R*.

##### (i) Shallow bud

For a shallow bud (***Figure 1–figure supplement 1B (i)***), which corresponds to buds smaller than a hemisphere, we can write that *A_c_ = A_m_* = 2*πR*^2^ (1 − cos *θ*), where 0 < *θ* < *π*/2 is the opening angle of the bud (see ***Figure 1–figure supplement 1B (i)***). In addition, *A_p_ = πρ*^2^ = *πR*^2^ sin^2^ *θ*; and *h = R*(1 − cos *θ*). Expressing these quantities as a function of the shape parameter, *η*, we obtain

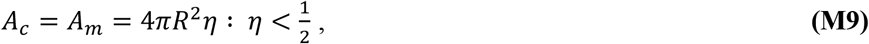

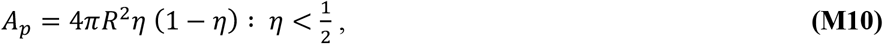

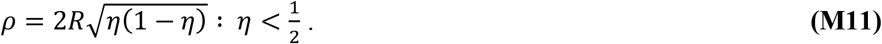

##### (ii) Deep bud

For a deep bud (***Figure 1–figure supplement 1B (ii)***), which corresponds to buds larger than a hemisphere, we can write that *A_c_* = 2*πR*^2^ (1 − cos *θ*), where *π*/2 < *θ < π*. In addition, *A_m_ = πR*^2^ (1 + (1 − cos *θ*)^2^); *A_p_ = πR*^2^; and *h = R*(1 − cos *θ*). Expressing these quantities as a function of the shape parameter, *η*, which in this case ranges between ½<*η* < 1, we obtain

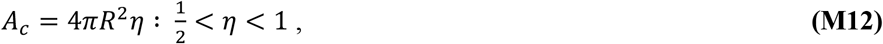

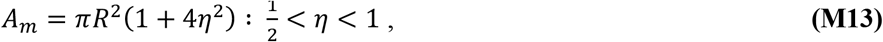

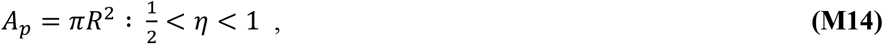

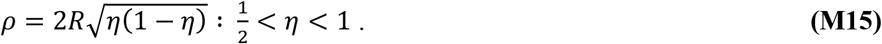

##### (iii) Pearled intermediate

A pearled intermediate corresponds to carriers form by an incomplete bud with opening angle 0 *< θ < π*, connected via a narrow connection with *n* complete buds (***Figure 1–figure supplement 1B (iii)***). Here, we can write that *A_c_* = 2*πR*^2^ [2*n* + (1 − *cos θ*)], where 0 < *θ < π* and *n* ≥ 1. In addition, *A_m_ = πR*^2^ [4*n* + 1 + (1 − *cos θ*)^2^]; *A_p_ = πR*^2^; and *h = R*(2*n* + 1 − cos *θ*). Expressing these quantities as a function of the shape parameter, *η*, we obtain

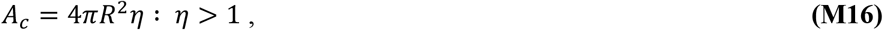

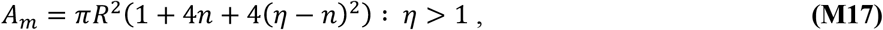

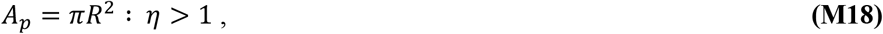

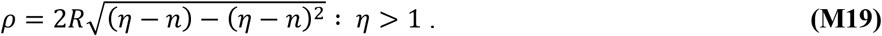

Putting together ***Equations (M9-19)***, we get:

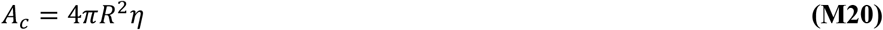

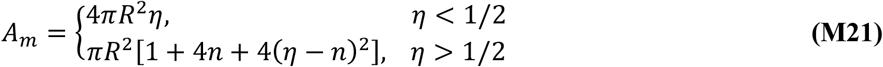

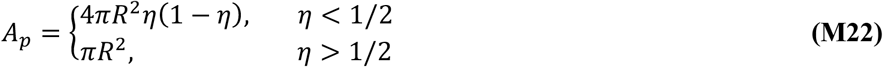

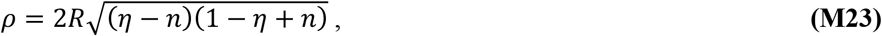

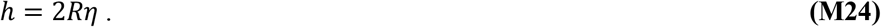

where *n* = [*η*] is the number of complete pearls, the brackets denoting the integer part operator. This allows us to express ***Equation (4)*** as

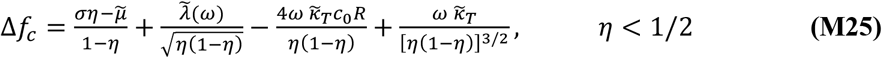

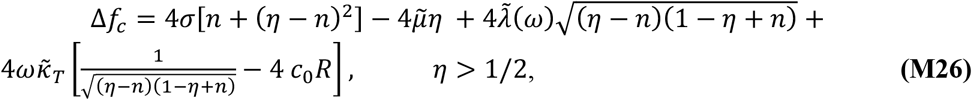

where 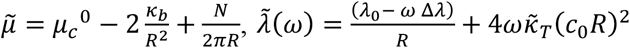, and 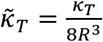 (***Equation (6)*** in the main text).

## ACKNOWLEDGEMENTS

We thank Javier Diego Íñiguez, Iván López-Montero, and members of the Garcia-Parajo lab for valuable discussions. M.F. Garcia-Parajo and V. Malhotra are Institució Catalana de Recerca i Estudis Avançats professors at ICFO-Institut de Ciencies Fotoniques and the Centre for Genomic Regulation (CRG), respectively. M. Chabanon, M.F. Garcia-Parajo and F. Campelo acknowledge support by the Spanish Ministry of Economy and Competitiveness (“Severo Ochoa” Programme for Centres of Excellence in R&D (SEV-2015-0522), BFU2015-73288-JIN, FIS2015-63550-R and FIS2017-89560-R), Fundacion Privada Cellex, Fundació Privada Mir-Puig, Generalitat de Catalunya through the CERCA program, ERC Advanced Grant NANO-MEMEC (GA 788546) and LaserLab 4 Europe (GA 654148). I. Raote and V. Malhotra acknowledge funding by grants from the Ministerio de Economía, Industria y Competitividad Plan Nacional (BFU2013-44188-P) and Consolider (CSD2009-00016); support of the Spanish Ministry of Economy and Competitiveness, through the Programmes “Centro de Excelencia Severo Ochoa 2013–2017” (SEV-2012–0208) and Maria de Maeztu Units of Excellence in R and D (MDM-2015–0502); and support of the CERCA Programme/Generalitat de Catalunya. I. Raote, M.F. Garcia-Parajo, V. Malhotra., and F. Campelo acknowledge initial support by a BIST Ignite Grant (eTANGO). I. Raote acknowledges support from the Spanish Ministry of Science, Innovation and Universities (IJCI-2017-34751). F. Campelo acknowledges support from the Ministerio de Ciencia, Innovación y Universidades (RYC-2017-22227). This work reflects only the authors’ views, and the EU Community is not liable for any use that may be made of the information contained therein. M. Arroyo and N. Walani acknowledge the support of the European Research Council (CoG-681434), and M. Arroyo that of the Generalitat de Catalunya (2017-SGR-1278 and ICREA Academia prize for excellence in research) and of the Spanish Ministry of Economy and Competitiveness, through the Severo Ochoa Programme (CEX2018-000797-S).

## SUPPLEMENTARY FIGURES

**Figure 1–figure supplement 1.**
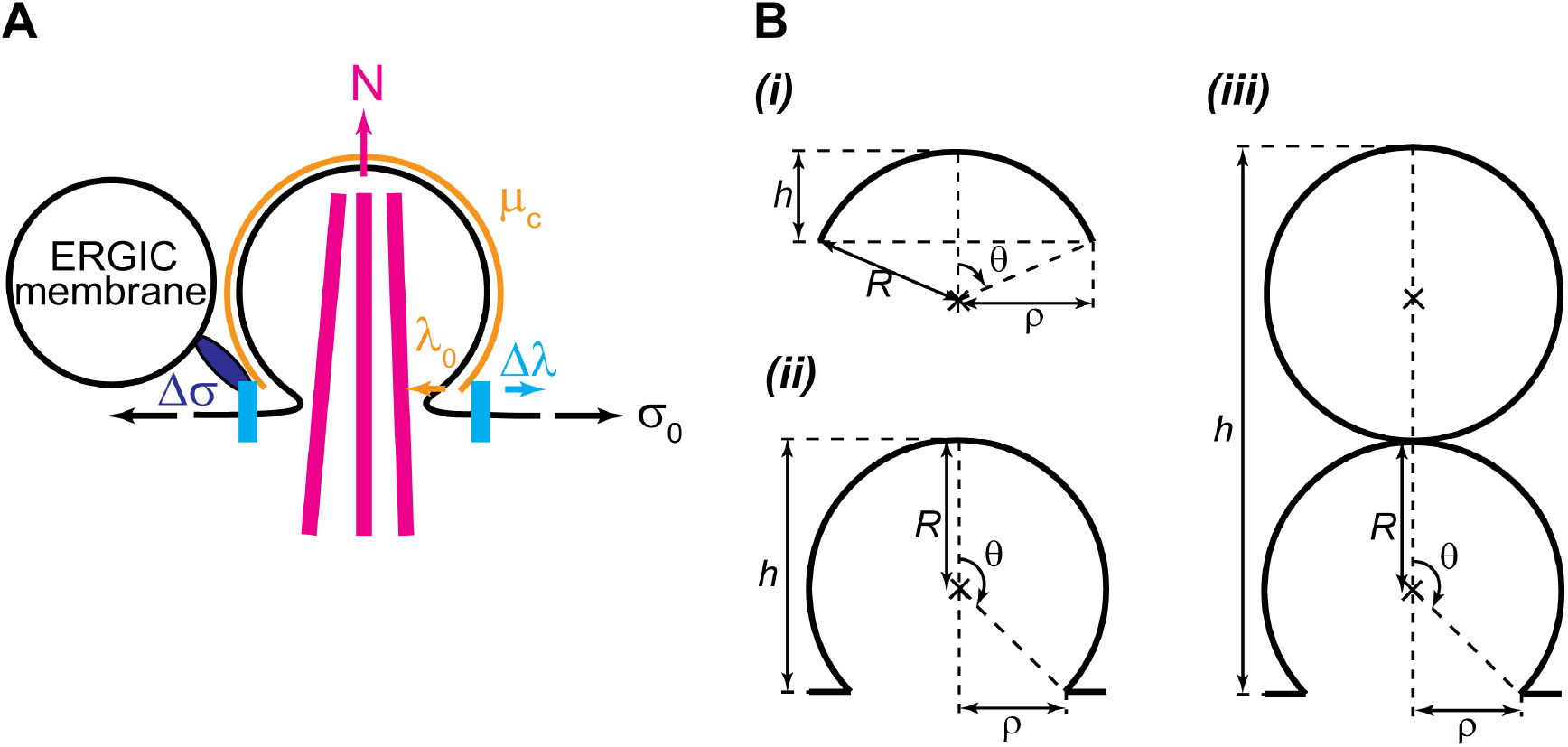
Geometry and physical forces in the transport intermediate generation model. **(A)** TANGO1 rings assembling on the ER membrane are depicted in light blue, accounting for a line tension reduction of the COPII coat, **Δ*λ***. The ER membrane is shown in black, associated with a tension, ***σ*_0_**. The COPII coat polymerizing on the membrane is depicted in orange, and accounts for a coat binding free energy (or chemical potential), ***μ_c_***, and a COPII coat line tension, ***λ*_0_**. Packaged procollagen rods are shown in magenta, which can (but not necessarily) contribute with a pushing normal force, ***N***. Finally, ERGIC53-containing membranes tethered to the export site through the NRZ complex (dark blue) can lead to a membrane tension reduction, **Δ*σ*. (B)** Scheme of the carrier geometry used for shallow buds *(i)*, deep buds *(ii)*; and pearled carriers *(iii)*. See Materials and Methods for the detailed description of the geometric parameters.

**Figure 2–figure supplement 1.**
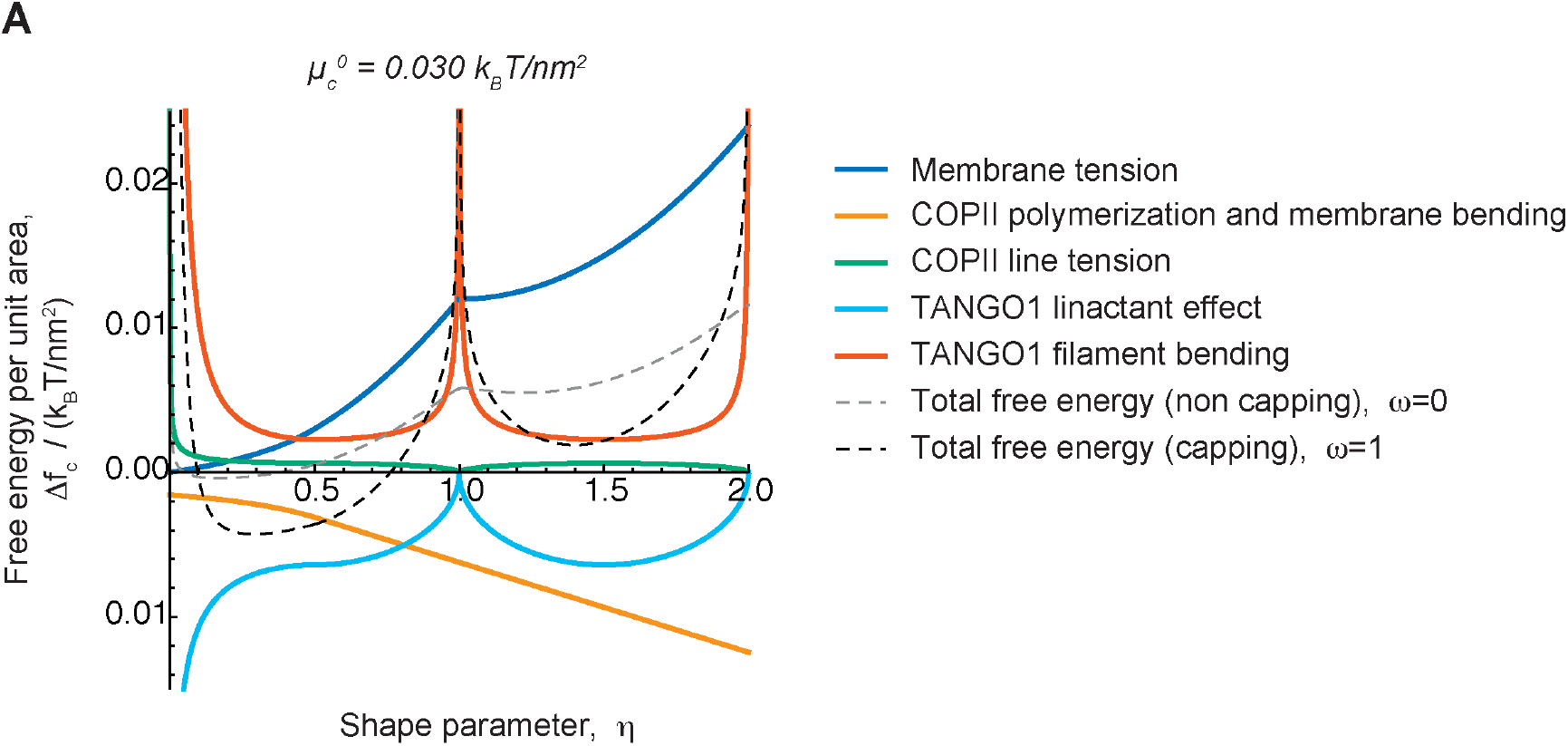
Different contributions to the free energy profile of a transport intermediate as a function of its shape and TANGO1 capping. **(A)** The free energy per unit area of the transport intermediate-TANGO1 system, Δ*f_c_*, plotted as a function of the shape parameter, *η*, for the COPII coat binding energy, 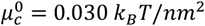, for both full COPII capping by TANGO1 *(*ω* = 1*, black, dashed curve) and full uncapping (*ω* = 0, gray, dashed curve), are shown, as in ***Figure 2A***. Each separate contribution to the total free energy per unit area (***Equations (4,5)***) is individually represented: the contributions of *(i)* membrane tension (dark blue); *(ii)* COPII polymerization and membrane bending (orange); *(iii)* COPII line energy (green); *(iv)* TANGO1-mediated reduction of the COPII line energy (light blue); and *(v)* TANGO1 filament bending energy (vermillion). Plots *(i–iii)* correspond to the free energies described by ***Equation (1)***; plot *(iv)* corresponds to the free energy described by ***Equation (3)***; and plots *(v)* corresponds to the free energy described by ***Equation (2)***. We considered no applied force, *N* = 0, and the rest of the elastic parameters used for the calculations are specified in ***Table 1***.

**Figure 5–figure supplement 1.**
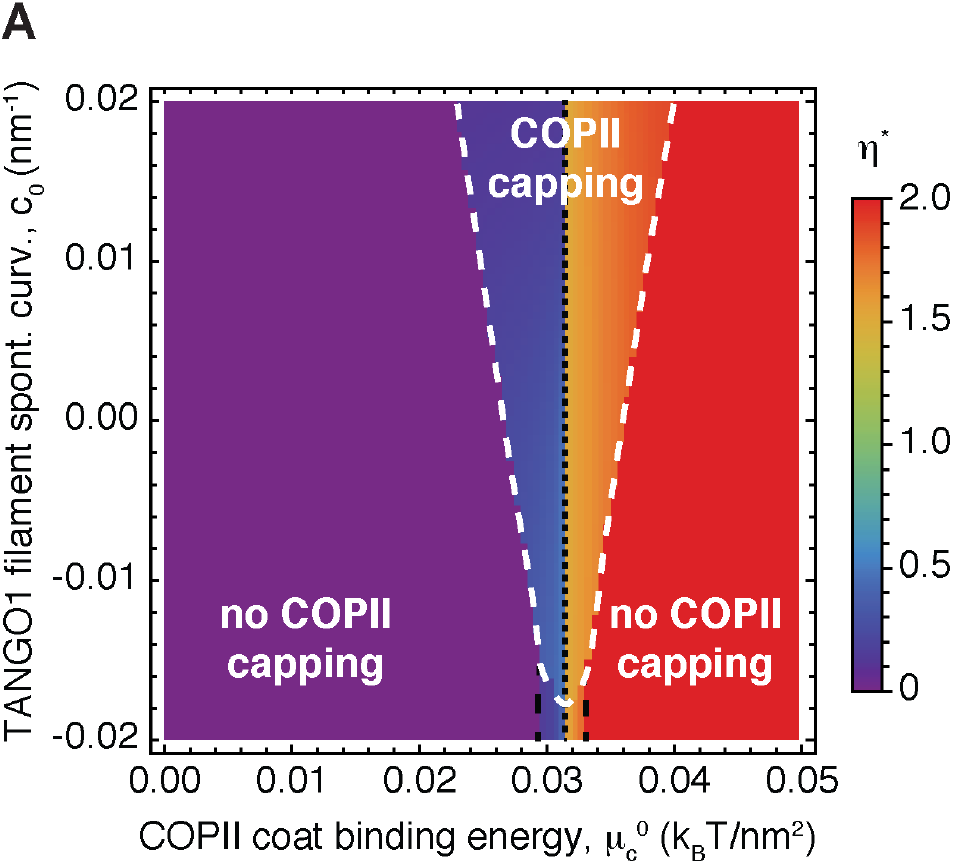
Shape diagram of the transport intermediate as a function of the TANGO1 filament spontaneous curvature. **(A)** Two-dimensional shape diagram indicating the shape of minimal elastic energy, represented by the optimal shape parameter, *η*^*^ (color-coded), and the state of TANGO1 capping (capping/uncapping transitions marked by thick, dashed, white lines). The shape diagram is plotted as a function of the COPII coat binding energy, 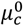, and of the TANGO1 filament spontaneous curvature, *c*_0_. There is no applied force (*N* = 0) and the rest of the elastic parameters used for all the calculations are listed in ***Table 1***.

**Figure 6–figure supplement 1.**
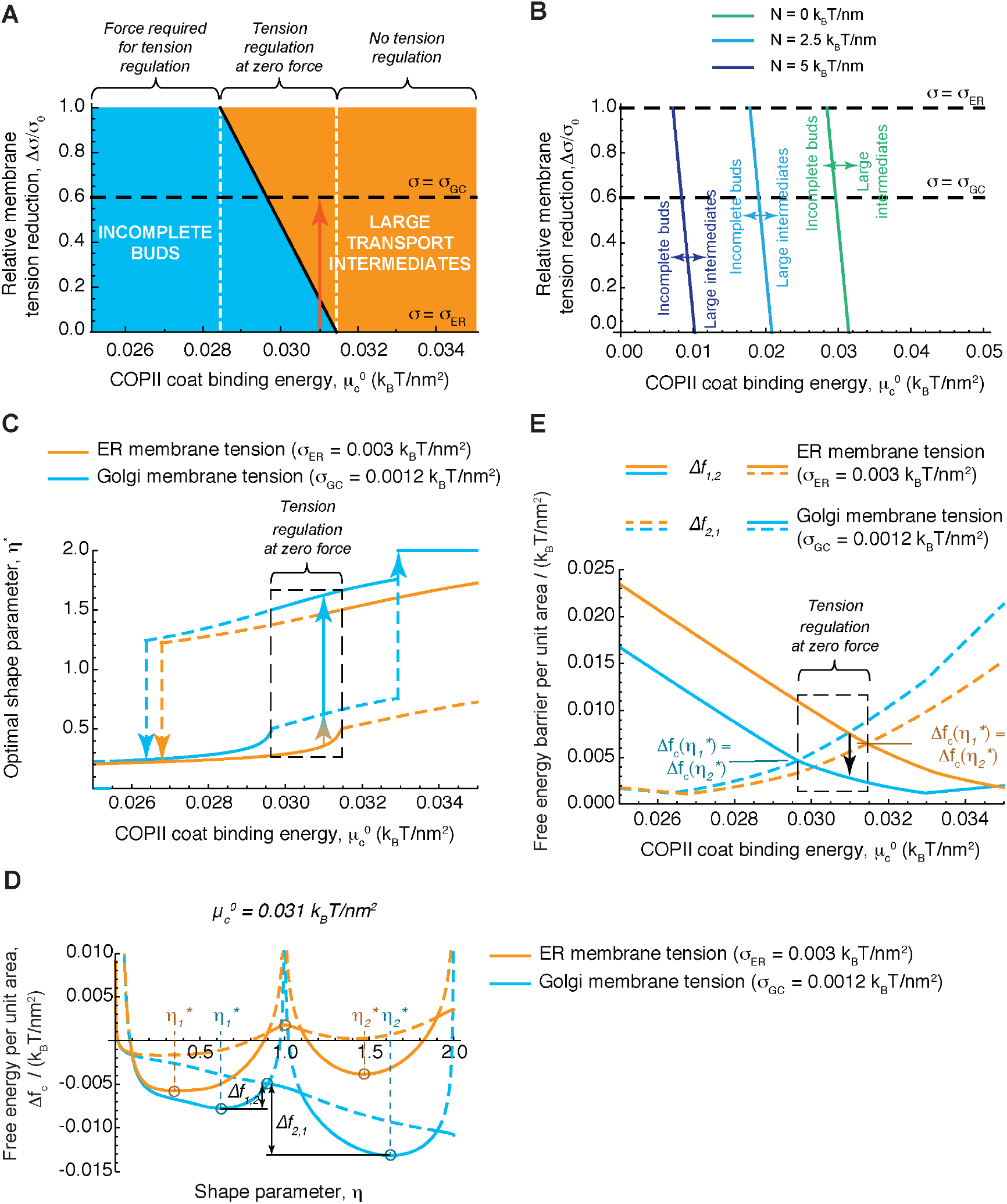
Membrane tension regulation by TANGO1 can mediate procollagen export. **(A)** Shape diagram showing the relative membrane tension reduction, Δ*σ/σ*_0_, as a function of the COPII coat binding energy, 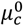, at zero applied force (*N* = 0). The corresponding shapes are indicated as incomplete buds (including both flat membranes and shallow buds), shown in blue, or large transport intermediates, in orange. Three distinct regions are indicated in the diagram, as in ***Figure 4A***, separated by vertical, dashed, white lines. The central region corresponds to the region of the parameter space where formation of large intermediates from incomplete buds can be triggered by a reduction in membrane tension. An example of such transitions, corresponding to relative tension reductions of 60% (from the measured ER tension, *σ_ER_*, to the measured Golgi membrane tension, *σ_GC_*, is indicated by a red arrow. The right and left regions show no possibility of large intermediate formation mediated by membrane tension regulation. In the left region, such regulation is still possible provided a force is pulling on the bud (see text for details). **(B)** Shape diagram as in **(A)** for different values of the applied force, *N* (see legend). The corresponding shapes are indicated as incomplete buds (including both flat membranes and shallow buds), to the left of the transition lines, or large transport intermediates, to the right of the transition lines. To illustrate how membrane tension reduction can trigger carrier elongation at different forces, the lines corresponding to measured ER tension, *σ_ER_*, and Golgi membrane tension, *σ_GC_*, are indicated by black, dashed lines. The right and left regions show no possibility of large intermediate formation mediated by membrane tension regulation. In the left region, such regulation is still possible provided a force is pulling on the bud (see text for details). **(C)** The optimal shape parameter, *η*^*^, is plotted as a function of the COPII coat binding energy, 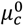, for two values of the membrane tension: *σ = σ_ER_* (orange curves), and *σ = σ_GC_* (blue curves). Globally stable shapes, for each case, are indicated by solid lines, whereas dashed lines represent locally, metastable shapes. Within the tension regulation at zero force region of the diagram (marked by the dashed, black box), a local, transient reduction in the membrane tension can lead to the growth of the transport intermediate (marked by the orange to blue faded arrows). **(D)** The free energy per unit area of the transport intermediate-TANGO1 system, Δ*f_c_*, is plotted as a function of the shape parameter, *η*, for the COPII coat binding energy, 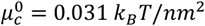, for both full COPII capping by TANGO1 (*ω* = 1) and full uncapping (*ω* = 0), and for two values of the membrane tension: *σ = σ_ER_* (orange curves); and *σ = σ_GC_* (blue curves). For each value of the shape parameter, *η*, the locally stable state of TANGO1 capping/unapping (lower free energy) is represented by the corresponding solid curve, whereas dashed curves indicate unstable states (higher free energy). The locally stable shapes, denoted by 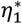 and 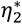 are indicated, as well as the free energy barriers,Δ*f*_1,2_, and Δ*f*_2,1_. **(E)** The free energy barriers separating the incomplete bud from the large intermediate morphologies, Δ*f*_1,2_ (solid lines), and Δ*f*_2,1_ (dashed lines) (defined in **(D)**), are plotted as a function of the COPII coat binding energy, *μ”*, for two values of the membrane tension: *σ = σ_ER_* (orange curves), and *σ = σ_GC_* (blue curves). Transition zone where the two energy barriers are of the same height, and hence both incomplete buds and large transport intermediates have the same free energy, are marked by a dashed, black box (tension regulation at zero force region). An arrow illustrates how decreasing the membrane tension leads to a reduction of the energy barrier for growth of the transport intermediate. Unless otherwise specified, the elastic parameters used for all the calculations shown are listed in ***Table 1***.

**Movie 1.**
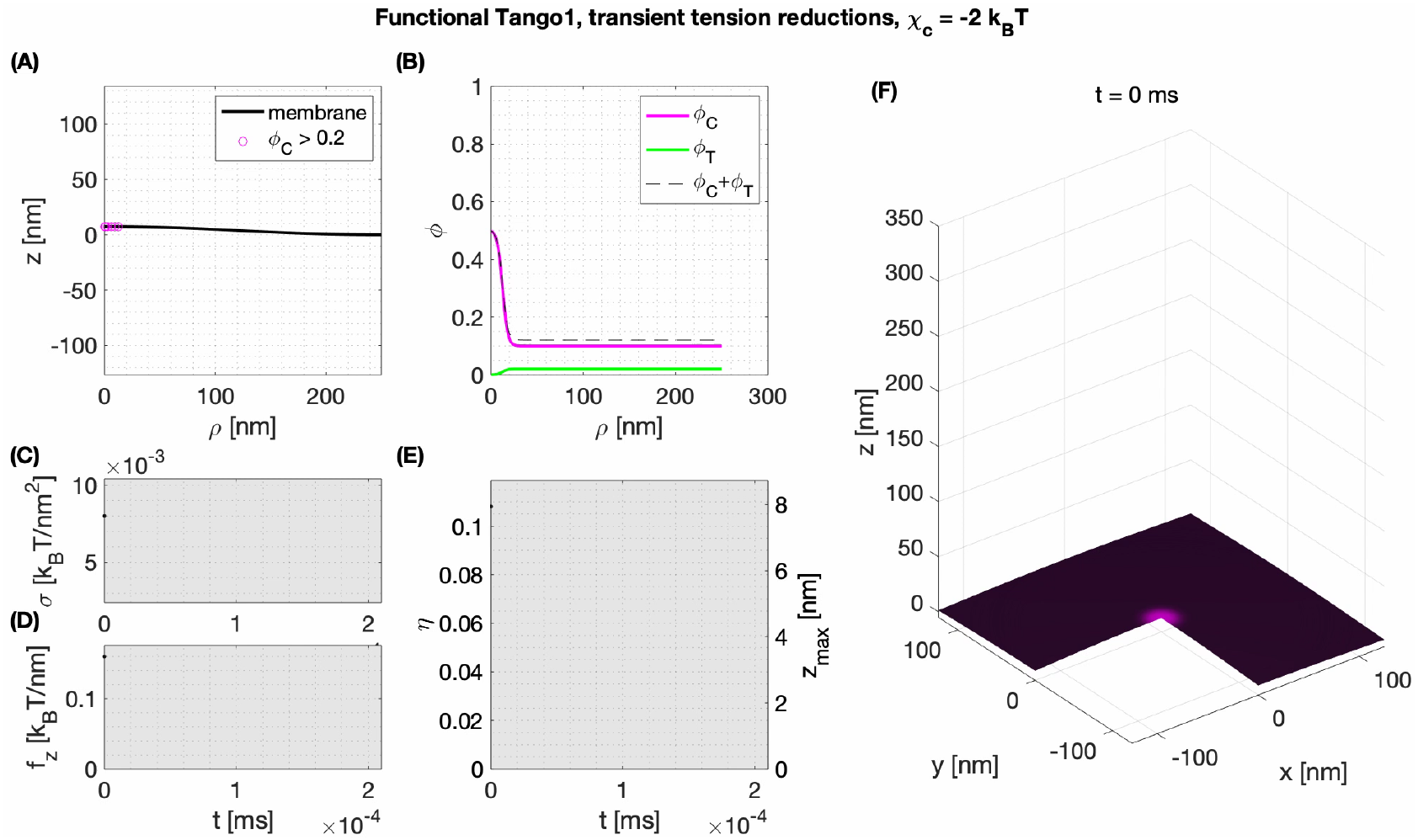
Membrane tension regulation by TANGO1 enables the formation of large procollagen-containing transport intermediates. Dynamics of an elongated COPII-coated transport intermediate formation with functional TANGO1. Membrane tension is transiently reduced from ***σ* = 0.008 *k_B_T/nm*^2^** to ***σ* = 0.004 *k_B_T/nm*^2^** and back ***σ* = 0.008 *k_B_T/nm*^2^** to allow the transition from a stable shallow bud to a stable “key-hole” shape, and then again to a stable “unduloid” shape of about 300 nm long, the typical size of a procollagen molecule. To mimic the presence of procollagen in the bud, a minimal neck radius if set at ***p_col_* = 7.5 *nm*** (see ***Appendix 1*** for details). **(A)** Shape of the membrane and distribution of proteins of more than 20% surface coverage. **(B)** Distribution of COPII and TANGO1 surface coverage as a function of the radius ***ρ*. (C)** Imposed membrane tension. **(D)** Imposed vertical point force, ***f_z_***, at ***ρ* = 0**. The value of ***f_z_*** is nonzero only during the initial COPII nucleation phase marked in gray. **(E)** Shape factor and bud height as a function of time. **(F)** Three-dimensional reconstitution of the transport intermediate and distribution of COPII (magenta) and TANGO1 (green).

## Appendix 1: Computational dynamic model of TANGO1-mediated bulky cargo export

In this Appendix we derive a dynamic model of a lipid bilayer whose spontaneous curvature is dictated by the diffusion and interaction of two membrane bound-species. We specialized this model to COPII and TANGO1, show that simulation results recapitulate key features of experimental observations, and use the model to propose biophysical mechanisms at the origin of transport intermediate formation for procollagen export from the ER.

The proposed model extends the work of [15] to account for a second membrane-bound species and apply it to TANGO1-COPII complex assembly. We therefore focus our exposé on this novel aspect, and direct readers interested in the detailed underlying theory to [2, 9, 14, 15].

### 1 Onsager’s variational approach: energetics, dissipation and power input

Our modeling approach is based on Onsager’s variational formalism of dissipative dynamics [2, 9]. The fundamental principle consists in describing the time evolution of the system through a minimization process of energy released, energy dissipated, and energy exchanged by the system. In other words, if 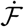 is the rate of change of free energy, 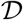 is the total dissipation potential, and 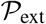 is the power input, the rate of change of the system can be obtained by minimizing at each time point the Rayleighian functional

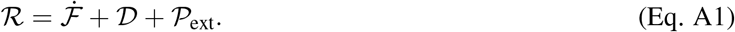

In this section, we define the energetic contributions to each of these quantities.

#### 1.1 Free energy

We consider a lipid bilayer as a material surface parametrized by **r**(*θ^α^, t*), where (*θ*^1^,*θ*^2^) are the Lagrangian surface coordinates, and *t* is the time. The state variable associated with the mechanical energy of the system is **r**, while the state variables associated with the chemical energy of the systems are the local area fractions of COPII and TANGO1, *ϕ_c_* and *ϕ_t_*, respectively. Note that these latter are bounded and should satisfy *ϕ_c_ > 0, ϕ_t_* > 0, and 0 ≤ *ϕ_c_ + ϕ_t_* ≤ 1.

##### 1.1.1 Mechanical energy

The bending energy of the membrane is described by the classical Helfrich model with spontaneous curvature [3, 4, 8]

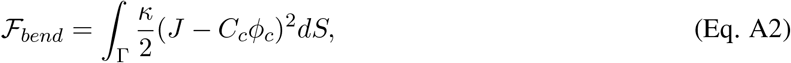

where *κ* is the bending modulus, *J* is the total curvature (twice the mean curvature, *H*) of the membrane, and the integration is performed over the entire membrane patch, Γ. Here we consider a local spontaneous curvature *C_c_ϕ_c_* resulting from the presence of COPII complexes. We assume a linear dependence of the spontaneous curvature on COPII local coverage, with *C_c_* = 2/*R_c_* being the maximum spontaneous curvature induced by a full coverage of COPII (*ϕ_c_* = 1) with preferred radius of curvature *R_c_*.

**Figure A1:**
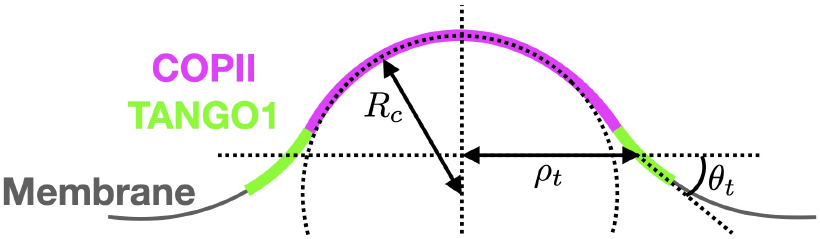
Schematic of a shallow COPII/TANGO1 bud and definition of the geometrical parameters favored by the proteins.

The total curvature *J* is a function of the state variable **r**. Briefly, from standard differential geometry [5, 6] we have that the tangent vectors at each point of the membrane are **g**_*α*_ = *∂***r**/*∂θ^α^*. They define the natural basis of the tangent space, from which the covariant components of the metric tensor are obtained *g_αβ_* = **g**_*α*_ · **g**_β_. Additionally the unit normal to the surface is 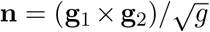, where *g* = det(*g_αβ_*). From these definitions, one gets the components of the second fundamental form *k_αβ_* = **n** · *∂***g**_*α*_/*∂θ^α^*, whose invariants are the total curvature *J* = tr **k** = *k_αβ_g^αβ^*, and the Gaussian curvature *K* = det **k** = *k_αβ_g^βγ^*. Here *g^βγ^* are the components of the inverse of the metric tensor, obtained from 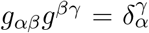. Note that for simplicity, we have neglected the contribution of the Gaussian curvature to the bending energy in (Eq. A2).

Based on our experimental observations [11], TANGO1 proteins are assumed to favor a ring-like conformation with a specific radius of curvature *ρ_t_* and a preferred angle with the plane of the ring *θ_t_* (see Fig. A1). As detailed later, we will restrict the model to axisymmetric shapes. Therefore for clarity, here we directly write an axisymmetric expression of the functional for the TANGO1 ring stiffness as

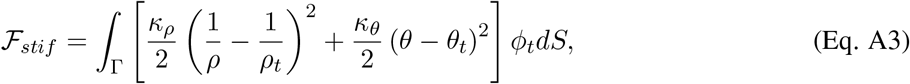

where *κ_ρ_* and *κ_θ_* are, respectively, the stiffness coefficients associated with the ring curvature 1/*ρ* and angle with the membrane *θ*.

##### 1.1.2 Chemical energy

We consider two distinct membrane-bound species representing COPII and TANGO1 proteins. They are described by continuous surface fractions *ϕ_c_* and *ϕ_t_*, respectively. We consider the entropic mixing energy of the two proteins to be represented by a Flory– Huggins type energy such as

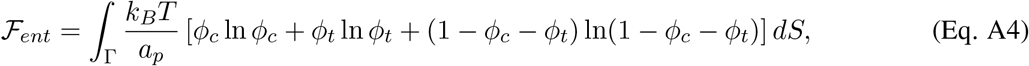

where *a_p_* is the molecular area of the proteins, assumed for simplicity to be identical for both proteins. The self-interaction and line energy of COPII proteins are

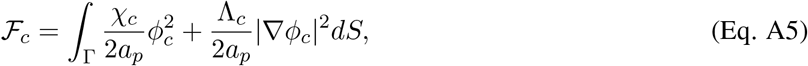

where *χ_c_* is negative for attractive interactions. The parameter Λ_*c*_ ensures a length-scale associated with the interface of the COPII domain: the spatial gradient of *ϕ_c_* is smoother for large values of Λ_*c*_.

Similarly, we write the self-interaction and line energy of TANGO1 proteins as

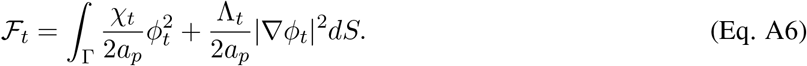

Finally, the interactions between COPII and TANGO1 proteins are

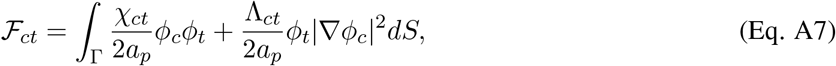

where the first term represents the affinity between the two proteins, and the second term represents the affinity of TANGO1 for the COPII coat boundary. Combining the second term of (Eq. A7) with the second term of (Eq. A5), one can see that TANGO1 modulates the interfacial energy of COPII with an effective parameter Λ_*c*_ + Λ_*ct*_ *ϕ_t_*.

The total free energy is the sum of the protein and membrane contributions:

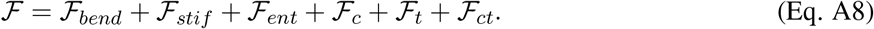

#### 1.2 Dissipation Mechanisms

Dissipation of mechanical energy occurs through in-plane shear stress of the lipid bilayer as it dynamically deforms under the action of a Lagrangian velocity of the membrane **V** = *∂***r**/*∂t* = **v**+*υ_n_***n**, where **v** and *υ_n_* are its tangential and normal components respectively. This membrane “flow” results in the time evolution metric tensor, whose time derivative in Lagrangian setting is the rate-of-deformation tensor of the surface [14]

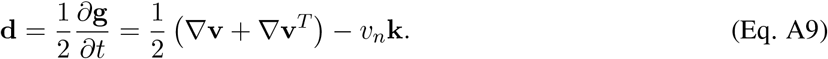

The first term, involving the membrane tangential velocity **v** represents the contribution of the tangential flow to the membrane deformation. The last term, involving the normal velocity *υ_n_*, accounts for the shape change of the membrane. The rate of change of local area is tr(d) = **▽ · v** − *υ_n_J*, which is zero for an inextensible membrane. We consider the lipid bilayer to be in a fluid phase that can be approximated by an interfacial viscous Newtonian fluid [1]. Neglecting intermonolayer friction, and assuming membrane inextensibility, the dissipation potential by in-plane shear stress takes the form

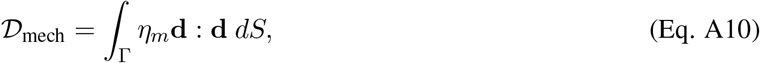

where *η_m_* is the in-plane viscosity of the lipid bilayer.

Dissipation of chemical energy occurs by protein diffusion along the membrane surface, described by the species diffusive velocities relative to the Lagrangian coordinates, *w_c_* and *w_t_* for COPII and TANGO1, respectively. Assuming for simplicity that the two proteins have the same molecular drag coefficient and surface area ap, the chemical dissipative potential of the system is

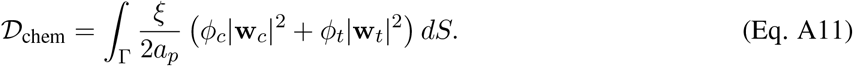

Given that the typical length scale of our system is well below the Saffman-Delbrück length scale (~ 1-10 *μ*m), we can safely neglect dissipation arising from the friction between the membrane and the cytosol [1]. The total dissipation potential of the system is 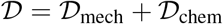.

#### 1.3 Power supplied

Mechanical power can only be supplied to our system through the boundary of the membrane patch *∂*Γ in the form of edge tractions and moments. Defining ***τ*** as the unit tangent vector along *∂*Γ so that ***ν = τ* × n**, the boundary tractions and moment power inputs are

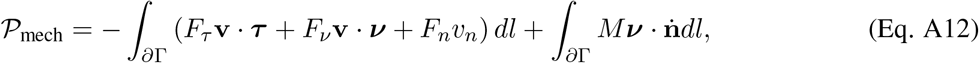

where *F_τ_, F_ν_* and *F_n_* are the traction components at the boundary, *M* is the bending moment per unit length, and n is the material time derivative of the surface normal.

In this model, we assume that all proteins are membrane bound and provided at the boundary of the domain by a protein reservoir of fixed chemical potential. The chemical power supply is therefore written

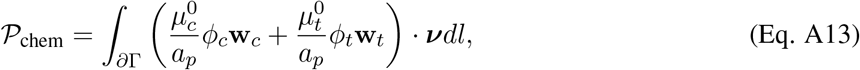

where 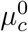 and 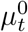 are the fixed boundary chemical potentials of COPII and TANGO1, respectively. The total power supplied to the system is 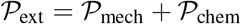.

### 2 Governing Dynamics

#### 2.1 Protein surface transport

Based on the definitions of the free energies, we define the energy density *W* as 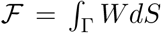. The chemical potentials of the two species can be written as *μ_i_ = a_p_* (*W_ϕ_i__* − ▽ · *W*_▽_*ϕ_i_*__), with *i* = {*c, t*}[15]. Here *W_ϕ_i__* and *W*_▽_*ϕ_i_*__ are the partial derivatives of W with respect to *ϕ_i_* and ▽_*ϕ_i_*_ respectively. The chemical potentials for each species are therefore

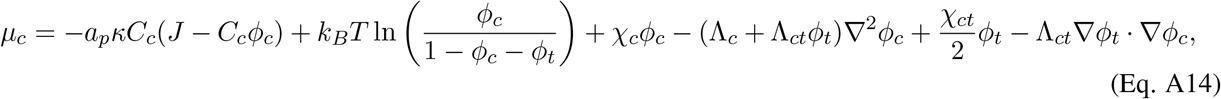

and

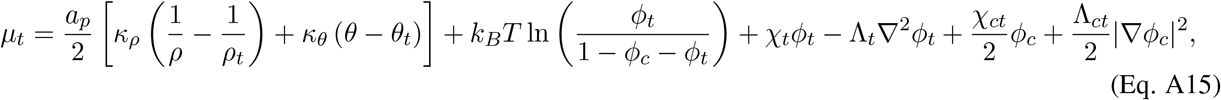

respectively.

The diffusive velocity of the species *i* can be expressed as a function of the species chemical potential *μ_i_* by minimizing the Rayleighian with respect to w_*i*_, giving w_*i*_ = −▽_*μ_i_*_/*ξ* [15]. The strong form of the transport equations for the proteins on an incompressible surface is therefore

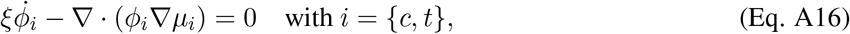

with the diffusive flux for each species given by

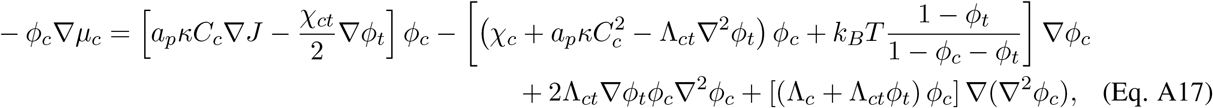

and

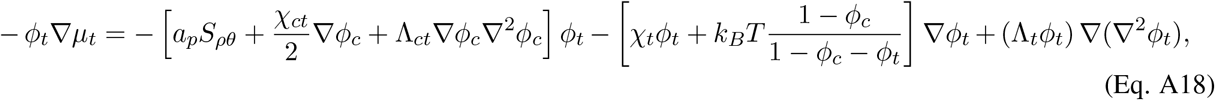

respectively. Here we defined *S_ρθ_ = κ_ρ_* (1/*ρ* − 1/*ρ_t_*)▽(1/*ρ*) + *κ_θ_*(*θ − θ_t_*)▽_*θ*_ and ▽^2^ is the surface Laplacian. These expressions highlight that COPII transport explicitly depends on the membrane curvature through the ▽_*J*_ term, while TANGO1 transport explicitly depends on the ring radius and its angle with the membrane through *S_ρθ_*.

#### 2.2 Membrane dynamics

To enforce local membrane incompressibility we introduce a Lagrange multiplier field *σ* that can be interpreted as the membrane tension [10]. Consequently, we aim at minimizing the Lagrangian functional

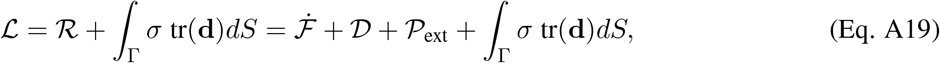

where 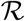 is the Rayleighian defined in (Eq. A1). Following Tozzi et al. [15], the dissipation rate of the free energy 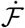 and the local area constraints can be expressed as functionals of the rate variables (**w, v**, *υ_n_*). The governing equations for the membrane mechanics are obtained by minimizing (Eq. A19) with respect to (**w, v**, *υ_n_*), and maximizing it with respect to *σ*. In the case of *W = W_bend_*, this results in the well-known shape equation and incompressibility condition [13, 17]. A full analysis of the governing equation for the total energy density of the COPII/TANGO1 system is out of the scope of this paper. For practical purposes, in what follows we proceed to a numerical minimization of (Eq. A19).

### 3 Model implementation

#### 3.1 Axisymmetric parametrization and numerical scheme

Details of the model implementation for a single species can be found in [15]. Briefly, we formulate the model in axisymmetric coordinates so that each material point of the membrane is expresses in terms of the distance to the axis of symmetry *ρ*(*u, t*) and of the axis of symmetry coordinate *z*(*u, t*). Here *u* ∈ [0, 1] is the Lagrangian coordinate along the membrane arclength, and *t* is time. The Dirichlet boundary conditions in the axisymmetric system take the form

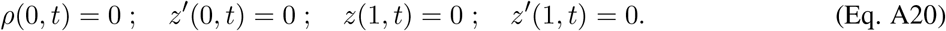

The fixed chemical potentials at the open boundary are ensured by

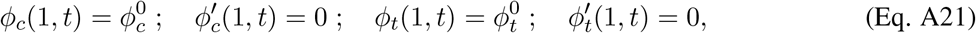

where 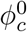 and 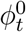 are the imposed protein densities of COPII and TANGO1, respectively, at the open boundary, mimicking protein reservoirs far from the budding site.

To solve numerically the coupled chemo-mechanical system, we employ a staggered approach where at each time step, we first solve the protein density field for a given membrane shape, and then update the shape at fixed membrane density distribution. The state variables are discretized using B-splines with cubic B-spline basis functions. The chemical problem is solved with the finite element method using a backward Euler discretization in time of the protein transport equations (Eq. A16), and Newton’s method to solve the resulting non-linear system. The mechanical problem is solved for a given distribution of proteins by minimizing the incremental Lagrangian from (Eq. A19) with respect to space variables and maximizing with respect to the Lagrange multiplier.

#### 3.2 Simulation and analysis protocols

After a preliminary parameter analysis informed by the physics of the problem (see also the main text for a discussion on parameters), we chose the reference set of parameters given in Table A1. Except stated otherwise, all results are obtained for these parameter values. All computations are done on an initially flat membrane patch of 250 nm radius.

**Table A1:** Model parameters

| Symbol | Parameter | Value |
| --- | --- | --- |
| *Material parameters* |  |  |
| $\kappa$ | Membrane bending rigidity | $20 \text{ k}_B\text{T}$ |
| $R_c$ | Preferred radius of curvature of COPII | $35 \text{ nm}$ |
| $C_c$ | COPII spontaneous curvature | $2/R_c \text{ nm}^{-1}$ |
| $\rho_t$ | TANGO1 preferred radius of curvature | $45 \text{ nm}$ |
| $\theta_t$ | TANGO1 preferred angle of curvature | $\pi/4$ |
| $\kappa_\rho$ | TANGO1 ring radius rigidity | $120 \text{ k}_B\text{T}$ |
| $\kappa_\theta$ | TANGO1 ring angle rigidity | $6.4 \times 10^{-4} \text{ k}_B\text{T nm}^{-2}$ |
| $a_p$ | Characteristic area of a protein | $100 \text{ nm}^2$ |
| $\chi_c$ | COPII self-interaction coefficient | $-2 \text{ k}_B\text{T}$ |
| $\chi_t$ | TANGO1 self-interaction coefficient | $-1.2 \text{ k}_B\text{T}$ |
| $\chi_{ct}$ | Affinity coefficient between COPII and TANGO 1 | $-0.4 \text{ k}_B\text{T}$ |
| $\Lambda_c$ | COPII interfacial coefficient | $2 \text{ k}_B\text{T}$ |
| $\Lambda_t$ | TANGO1 interfacial coefficient | $0.4 \text{ k}_B\text{T}$ |
| $\Lambda_{ct}$ | Coupling interfacial coefficient | $0.4 \text{ k}_B\text{T}$ |
| $\eta_m$ | Membrane viscosity | $5 \times 10^{-9} \text{ N s m}^{-1}$ |
| $\xi$ | Molecular drag coefficient ($= 2\pi\eta_m$) | $3.14 \times 10^{-8} \text{ N s m}^{-1}$ |
| *Model constraints* |  |  |
| $\phi_c^0$ | COPII protein surface fraction at the open boundary | $0.1$ |
| $\phi_t^0$ | TANGO1 protein surface fraction at the open boundary | $0.02$ |
| $\sigma$ | Membrane tension | $0.004 - 0.0096 \text{ k}_B\text{T/nm}^2$ |
| $f_z$ | Vertical axial force (applied for nucleation) | $0 - 0.56 \text{ k}_B\text{T/nm}$ |
| $\rho_{\text{col}}$ | Minimal neck radius in presence of procollagen molecules | $7.5 \text{ nm}$ |

##### Stable equilibrium

We assume that an equilibrium shape is reached if the maximum displacement between two time-steps of each material points is below 5 × 10^−3^ nm for more than ten time-steps in a row over a cumulative time larger than 1ms.

##### Shape parameter

To facilitate a quantitative comparison between the end-states of the system obtained for different sets of parameters, we use the shape parameter, which is essentially the maximum height of the bud normalized by the preferred diameter of curvature of COPII *η* = zmax/2R_*c*_ (see also main text).

##### Coat nucleation

In our system, the flat membrane state is a locally (sometimes globally) stable equilibrium state. This means that a perturbation needs to be applied to initiate the nucleation of COPII coats. To ensure a reproducible perturbation protocol, we define a criteria for COPII coat nucleation such as the average surface density of COPII within a surface area *A*_nuc_ = *π*(25 nm)^2^ around the axis of symmetry must satisfy ∫_*A*_nuc__ *ϕ_c_ dS* > 0.75 *A*_nuc_. Starting from a flat membrane patch with homogeneous distribution of species, a small upward point force of *f_z_* =0.16 k_B_T/nm is applied at *ρ* = 0. The force induces membrane deformation and initiates the recruitment of COPII. If the force is sufficient to nucleate a COPII coat as defined above, the point force is set back to zero, and the system is free to evolve. Alternatively, if an equilibrium is reached but the nucleation criteria is not satisfied, we gradually increase *f_z_* until either a COPII coat is nucleated or until *f_z_* >0.56k_B_T/nm, in which case we assume the flat membrane to be the stable state. The up-ward point force is implemented within the arclength parametrization by setting *F_n_*(0, *t*) = *f_z_* in the power supply (Eq. A12).

##### Neck closure

In the cases where the transport carrier closes, no equilibrium is reached. Consequently, we assume that if the neck radius goes below 5 nm, we reach the small length scale limit of the continuum modeling approach, and assume neck closure. Because the neck closure event happens at different times after the neck snap-through, in order to facilitate the comparison of the carrier height in a systematic manner in Fig. A3, we take *η* at the minimum bud height after the neck snaps.

##### Prevention of neck closure by collagen molecules

In simulations where procollagen molecules prevent the total closure, an energy penalty ∫_Γ_ 10^−3^*κ*/(*ρ − ρ*_col_)^2^*dS* is imposed on the portion of the membrane *u* ∈ [0.1; 0.8] using a hyperbolic tangent.

### Results and discussion

**Figure A2:**
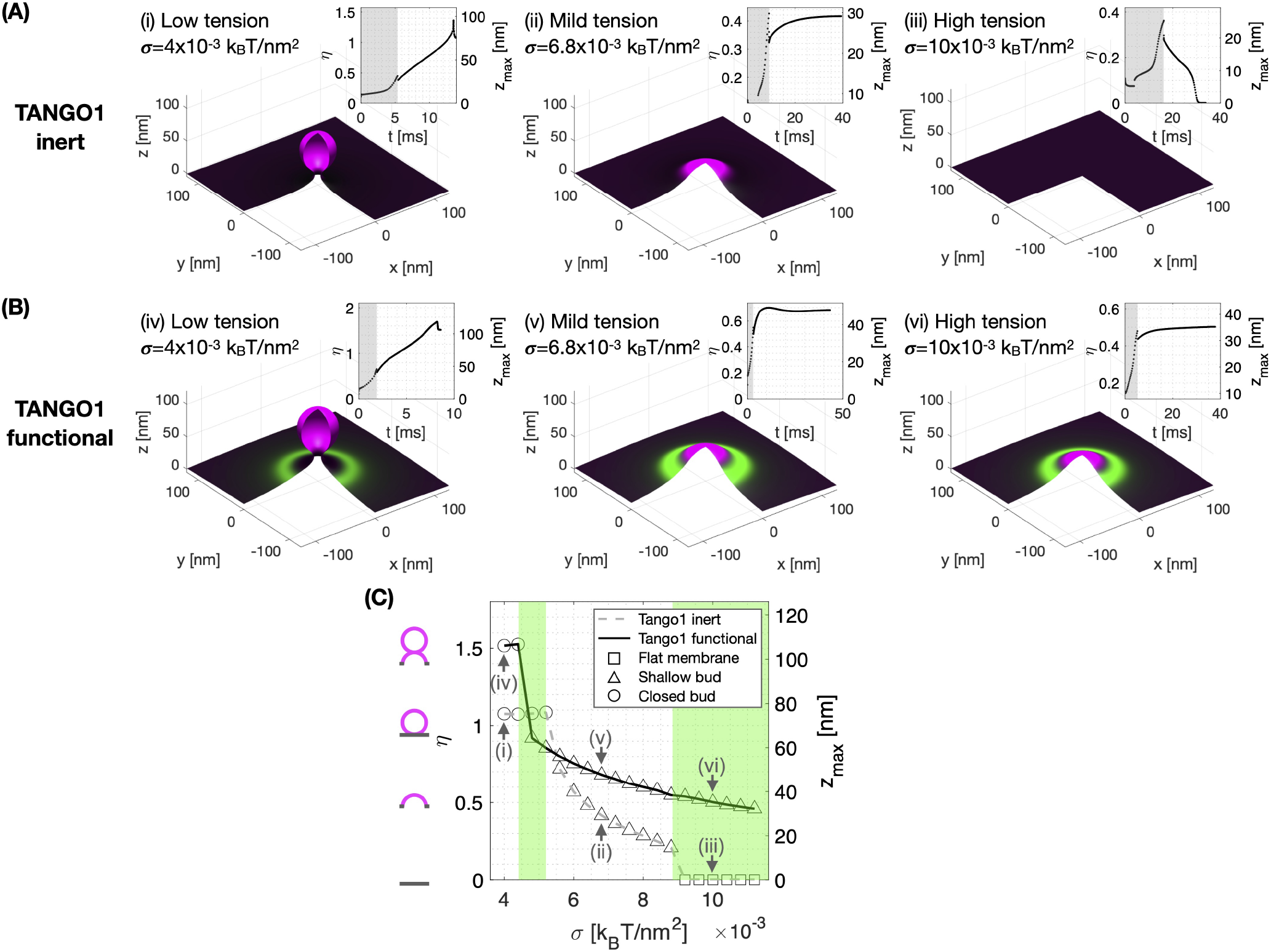
Computational results show that TANGO1 forms stable rings around COPII coats, and favors stable shallow buds over a larger range of membrane tension. (A, B) Final stable shapes at different membrane tensions for (A) inert TANGO1 but functional COPII and (B) functional TANGO1 and COPII. Inserts show the shape factor (*η* = *z*_max_/2R_*c*_) and height (zmax) as a function of time (gray regions indicate the perturbation stage where an incremental upward point force is applied to initiate the COPII coat nucleation). (C) Shape factor and bud height as a function of membrane tension. Each symbol represents the final state of a simulation with a different membrane tension (lines are guides for the eye). The final shape of the COPII coated transport intermediate is classified as stable closed spherical bud (circles), stable shallow bud (triangles), or stable flat membrane (squares). Green regions highlight tension regimes where TANGO1 mediates the regulation of the final membrane shape. All results obtained for parameters as given in Table A1, and *χ_c_* = −1.9 k_B_T. (See corresponding Supporting Movies 1-6 for a dynamic representation of the simulations).

**Figure A3:**
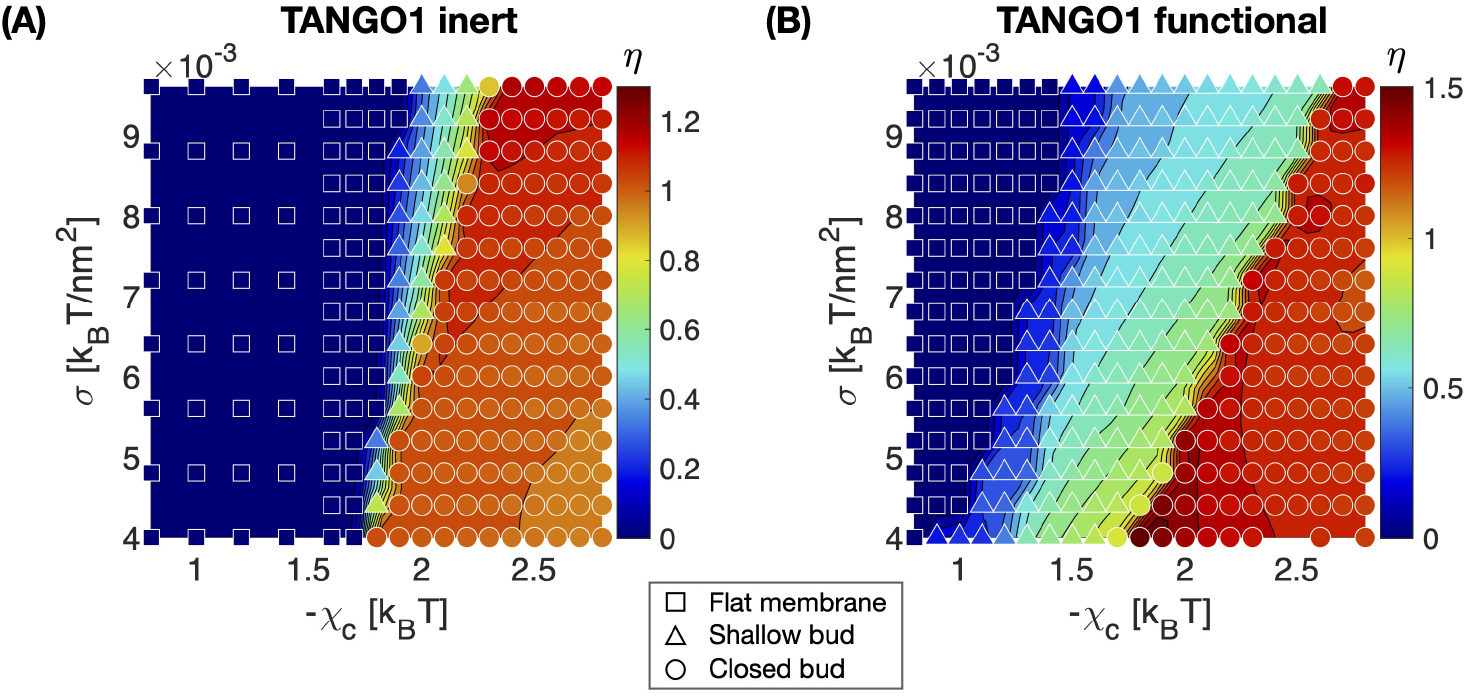
TANGO1 significantly widens the range of COPII self-interaction and membrane tension at which shallow buds are stable. Shape factor (color code) and final state (symbol) as a function of membrane tension (*σ*) and COPII self-interaction (*χ_c_*). (A) Inert TANGO1 but functional COPII. (B) Functional TANGO1 and COPII. Each symbol represents the final state of a dynamic simulation, while isocontours and colors are interpolated from the values of *η* at each symbol.

We first investigate the ability of COPII complex alone to generate spherical carriers. This corresponds to the control case where TANGO1 is treated as an inert species that only contributes to the entropic energy (*χ_t_, χ_ct_, κ_ρ_, κ_θ_* are set to zero). As shown in Fig. A2A, the final shape of the carrier depends on the membrane tension *σ*. At low membrane tension (Fig. A2A(i) and Supporting Movie 1), a COPII coat polymerizes and induces membrane curvature such that a spherical carrier forms and closes with a neck radius below the threshold of 5 nm. At intermediate and high membrane tensions, the COPII coats do not reach closure, and either stabilize a shallow state (Fig. A2A(ii) and Supporting Movie 2) or depolymerize and flatten out (Fig. A2A(iii) and Supporting Movie 3), respectively. As expected, the inert TANGO1 species does not play a role outside of entropically slowing down COPII dynamics. The result of our dynamic simulations with inert TANGO1 extend the conclusion of previous models obtained at equilibrium that highlighted the role of membrane tension as a regulator of bud formation [7, 12, 16].

In the presence of functional TANGO1, we observe the formation of a stable TANGO1 ring around the COPII coats (Fig. A2B and Supporting Movies 4-6). Importantly, we find that at high tension (Fig. A2B(vi)), TANGO1 prevents the membrane to flatten out by stabilizing the COPII coat in a stable shallow bud. Such TANGO1-assisted transport carrier regulation is better seen in Fig. A2C, where the bud height and final states are plotted as a function of the membrane tension for both inert and functional TANGO1 simulations. We find that functional TANGO1 not only increases the overall bud height compared to inert TANGO1, but also widens the range of tensions at which shallow buds are stable, both at high and intermediate tension regimes (green regions in Fig. A2C). This effect is critical at intermediate tensions corresponding to the transition from shallow to closed buds: the presence of functional TANGO1 lowers the minimum membrane tension required to close a bud, therefore indicating that TANGO1 could potentially play a role in tension-regulated transport intermediate formation.

To better understand the competition between membrane tension and COPII coat polymerization during transport carrier formation, we show in Fig. A3 the final carrier states and shape factor as a function of membrane tension (*σ*) and COPII self-interaction coefficient (*χ_c_*) for both inert and active TANGO1. We find that at low COPII self-interaction, the COPII coat either does not nucleate during the initial perturbation stage, or does nucleate but cannot provide enough bending energy to stabilize the membrane curvature against the membrane tension, thus resulting in a flat membrane (*η* = 0). At high COPII self-interaction, the nucleated coat successfully works against membrane tension to generate membrane curvature, generating a closed spherical bud. At intermediate COPII self-interaction, we find a transition region where the deformed membrane stabilizes as an open, shallow bud. Interestingly, the presence of functional TANGO1 dramatically widens the parameter space at which stable shallow buds are obtained. Importantly, the phase boundary at the transition from shallow to closed bud now spans a larger range of membrane tension and COPII self-interaction. Note that the COPII self-interaction parameter *χ_c_* is conceptually equivalent to *μ*^0^ in the equilibrium model presented in the main text. Therefore the numerical results shown in Fig. A3 are in a good qualitative agreement with the equilibrium model results (Fig. 4A in the main text). Despite the different assumptions underlying the computational dynamic and equilibrium models, their qualitative agreement highlights the relevance of their common physical mechanisms to TANGO1-mediated assembly of procollagen-containing transport intermediates.

These observations suggest that stable shallow COPII buds surrounded by TANGO1 rings can remain stable in the ER, possibly facilitating the recruitment and packaging of procollagens to ERES. Additionally, our results suggest that a shallow bud stabilized by TANGO1 could close into a spherical carrier if the membrane tension is transiently reduced. Such tension reduction hypothesis is supported experimentally by the recruitment of ERGIC53-containing membrane vesicles by TANGO1 rings through the NRZ tether complex [11].

In summary, we outlined here the fundamentals of a full dynamic model for lipid bilayers with two membranebound species (TANGO1 and COPII complexes, in particular). We applied this model to the study of how TANGO1 can modulate the formation of COPII-coated transport intermediates. Our results are in a good qualitative agreement with the results presented in the main text obtained by using an analytical equilibrium model. A full parameter exploration of this model is beyond the scope of this article. However, we expect that the dynamic model outlined here will be very valuable to test new hypotheses based on ongoing and future experimental results on this and other intriguing cellular processes.

## Supporting movie captions

Movies 1-6 are the dynamic simulations corresponding to the end states shown in pannels (i-vi) of Fig. A2 respectively.

Supporting movie 1: Dynamic of COPII bud formation with inert TANGO1 at low membrane tension (*σ* = 4 k_B_T/nm^2^) and intermediate COPII self interaction (*χ_c_* = −1.9 k_B_T). The final state is a closed bud. (A) Shape of the membrane and distribution of proteins of more than 20% surface coverage. (B) Distribution of COPII and TANGO1 surface coverage as a function of the radius *ρ*. (C) Imposed membrane tension. (D) Imposed vertical point force (*f_z_*) at *ρ* = 0. The value of *f_z_* is non-zero only during the initial COPII nucleation phase marked in gray. (E) Shape factor and bud height as a function of time. (E) Three-dimensional reconstitution of the bud and distribution of COPII (magenta) and TANGO1 (green).

Supporting movie 2: Dynamic of COPII bud formation with inert TANGO1 at intermediate membrane tension (*σ* = 6.8 k_B_T/nm^2^) and intermediate COPII self interaction (*χ_c_* = −1.9 k_B_T). The final state is a stable shallow bud. (A) Shape of the membrane and distribution of proteins of more than 20% surface coverage. (B) Distribution of COPII and TANGO1 surface coverage as a function of the radius *ρ*. (C) Imposed membrane tension. (D) Imposed vertical point force (*f_z_*) at *ρ* = 0. The value of *f_z_* is non-zero only during the initial COPII nucleation phase marked in gray. (E) Shape factor and bud height as a function of time. (E) Three-dimensional reconstitution of the bud and distribution of COPII (magenta) and TANGO1 (green).

Supporting movie 3: Dynamic of COPII bud formation with inert TANGO1 at high membrane tension (*σ* = 10 k_B_T/nm^2^) and intermediate COPII self interaction (*χ_c_* = −1.9 k_B_T). The final state is a stable flat membrane. (A) Shape of the membrane and distribution of proteins of more than 20% surface coverage. (B) Distribution of COPII and TANGO1 surface coverage as a function of the radius *ρ*. (C) Imposed membrane tension. (D) Imposed vertical point force (*f_z_*) at *ρ* = 0. The value of *f_z_* is non-zero only during the initial COPII nucleation phase marked in gray. (E) Shape factor and bud height as a function of time. (E) Three-dimensional reconstitution of the bud and distribution of COPII (magenta) and TANGO1 (green).

Supporting movie 4: Dynamic of COPII bud formation with functional TANGO1 at low membrane tension (*σ* = 4 k_B_T/nm^2^) and intermediate COPII self interaction (*χ_c_* = −1.9 k_B_T). The final state is a closed bud.(A) Shape of the membrane and distribution of proteins of more than 20% surface coverage. (B) Distribution of COPII and TANGO1 surface coverage as a function of the radius *ρ*. (C) Imposed membrane tension. (D) Imposed vertical point force (*f_z_*) at *ρ* = 0. The value of *f_z_* is non-zero only during the initial COPII nucleation phase marked in gray. (E) Shape factor and bud height as a function of time. (E) Three-dimensional reconstitution of the bud and distribution of COPII (magenta) and TANGO1 (green).

Supporting movie 5: Dynamic of COPII bud formation with functional TANGO1 at intermediate membrane tension (*σ* = 6.8 k_B_T/nm^2^) and intermediate COPII self interaction (*χ_c_* = −1.9 k_B_T). The final state is a stable shallow bud. (A) Shape of the membrane and distribution of proteins of more than 20% surface coverage. (B) Distribution of COPII and TANGO1 surface coverage as a function of the radius *ρ*. (C) Imposed membrane tension. (D) Imposed vertical point force (*f_z_*) at *ρ* = 0. The value of *f_z_* is non-zero only during the initial COPII nucleation phase marked in gray. (E) Shape factor and bud height as a function of time. (E) Three-dimensional reconstitution of the bud and distribution of COPII (magenta) and TANGO1 (green).

Supporting movie 6: Dynamic of COPII bud formation with functional TANGO1 at high membrane tension (*σ* = 10 k_B_T/nm^2^) and intermediate COPII self interaction (*χ_c_* = −1.9 k_B_T). The final state is a stable shallow bud. (A) Shape of the membrane and distribution of proteins of more than 20% surface coverage. (B) Distribution of COPII and TANGO1 surface coverage as a function of the radius *ρ*. (C) Imposed membrane tension. (D) Imposed vertical point force (*f_z_*) at *ρ* = 0. The value of *f_z_* is non-zero only during the initial COPII nucleation phase marked in gray. (E) Shape factor and bud height as a function of time. (E) Three-dimensional reconstitution of the bud and distribution of COPII (magenta) and TANGO1 (green).

